# Haplotype tagging sheds light on speciation between two divergent cryptic species of the brown algae *Ectocarpus*

**DOI:** 10.64898/2026.05.05.723031

**Authors:** Cécile Molinier, Lauric Reynes, Yingguang Frank Chan, Marek Kučka, Jérôme Coudret, Remy Luthringer, Fabian B. Haas, Akira F. Peters, Alejandro E. Montecinos, Marie-Laure Guillemin, Christophe Destombe, Susana M. Coelho, Myriam Valero, Agnieszka P. Lipinska

## Abstract

Understanding how genome-wide divergence translates to barriers to gene flow is a central question in speciation research, particularly in marine environments where interconnected habitats frequently accommodate cryptic diversity. Using linked-read whole-genome data from the brown alga *Ectocarpus*, we investigated genome-wide reproductive isolation between two cryptic brown algal species *E. siliculosus* and *E. crouaniorum* across replicated hybrid zones in Europe and Chile. We first uncover deep genetic structure within *E. crouaniorum*, revealing a more intricate species complex than previously recognized. Despite this complexity, we found parallel patterns of asymmetric introgression that emerged independently in Europe and Chile, where we detected both, F1 and low-level introgressed individuals. However, the specific lineage pairs involved in hybridization depended on local demographic history. Demographic modeling further indicated that reproductive isolation strengthens gradually with divergence time and differs markedly between geographic regions, with ongoing asymmetric gene flow even between lineage pairs where hybrids were not detected in the field. In all cases, the genomic landscape of hybridization is consistent with a polygenic, genome-wide barrier to gene flow rather than a few large-effect regions. Our results show that cryptic speciation in brown algae is driven by repeatable, regionally parallel architectures of genome-wide reproductive isolation which is shaped by ecological and demographic context.

## 1. INTRODUCTION

The process of speciation is tightly linked to the evolution of reproductive isolation (RI), which limits genetic exchange between diverging populations (Coyne & Orr, 2004; Westram et al., 2022). In the classical allopatric model, physical barriers aid speciation by restricting gene flow, which allows populations to accumulate genetic differences over time through drift or selection (Coyne & Orr, 2004). Given enough time, genome-wide differentiation can become fixed, leading to reproductive barriers that persist even after secondary contact (Coyne & Orr, 2004; Mayr, 1942). However, growing evidence suggests that speciation can also occur in the presence of gene flow and could be driven by divergent selection acting on specific traits despite the homogenizing effect of recombination (Feder et al., 2012; Nosil, 2008; Smadja & Butlin, 2011). In such cases, genomic divergence may be found at the localized "speciation islands," which govern crucial traits that maintain species boundaries (Feder et al., 2012; Via, 2012; Wolf & Ellegren, 2017; Wu, 2001). Alternatively, selection may also act simultaneously on multiple independent genomic regions resulting in “continents” of differentiation (Michel et al., 2010; Renaut et al., 2012; Sendell-Price et al., 2020). While these contrasting genomic patterns of speciation have been extensively studied in plants and animals (Le Moan et al., 2024; Nadeau et al., 2012; Renaut et al., 2012; Stankowski et al., 2019), other eukaryotic groups, such as marine algae, still remain largely unexplored. This leaves an open question of how genome-wide divergence is associated with barriers to gene flow in these organisms.

The marine environment presents a particularly interesting setting for studying speciation with gene flow. By contrast to terrestrial and freshwater habitats, where physical barriers can drive divergence, the open nature of the ocean facilitates the dispersal of spores, zygotes, propagules and drifting individuals over long distances (Brennan et al., 2014; Buchanan & Zuccarello, 2012; Fraser et al., 2009; McKenzie & Bellgrove, 2008; Palumbi, 1992). However, this paradigm of high gene flow in marine environments was first challenged by Palumbi in 1994 (Palumbi, 1994) and has since been supported by numerous population genetics studies (Bierne et al., 2003; Ravinet et al., 2021; Reynes et al., 2026). Part of the explanation likely lies in ecological factors. Niche specialization, host specificity in epiphytic algae, sexual behaviour and local adaptation can all impose strong selective pressures and help maintain species boundaries even in highly connected environments (Bierne et al., 2003; Peters et al., 2010; Solas et al., 2024; Solignac, 1981). In parallel, neutral processes like geographic distance and ocean current patterns further contribute to differentiation and isolation by facilitating genetic drift (Lebret et al., 2012; Thomas et al., 2015). Finally, global change can impact speciation dynamics through change in species distribution ranges. Previously isolated populations are brought back into contact, sometimes leading to hybridization which could further accelerate or hinder speciation (Hudson et al., 2021). Studying how past population history and local environmental factors interact to shape reproductive isolation is therefore essential for understanding the origin and maintenance of marine biodiversity.

Brown algae (Phaeophyceae) offer powerful systems to investigate these questions. They diverged from plants and animals over one billion years ago and evolved complex multicellularity independently (Baldauf, 2003; Cock et al., 2010). A defining feature of the majority of brown algae is their haploid-diploid life cycle, with alternation between multicellular free-living haploid gametophyte and diploid sporophyte generations (Cock et al., 2014; Müller, 1964) (Fig. 1A). Haploid gametophytes produce male and female gametes that fuse to form a diploid sporophyte, which subsequently undergoes meiosis to generate haploid meio-spores that develop into new gametophytes (Fig. 1A). This biphasic life history provides a unique opportunity to disentangle the role of selection acting on haploid and diploid phases in shaping reproductive barriers. In hybrid diploid sporophytes, recessive deleterious interactions may be hidden by allelic masking, whereas genomic incompatibilities in recombinant haploid gametophytes are immediately exposed to selection, potentially accelerating the evolution of reproductive barriers (Montecinos, Guillemin, et al., 2017; Nouhaud et al., 2020). However, structural barriers, such as chromosomal mispairing at meiosis, or genic incompatibilities during formation of spores can still result in F1 sterile sporophytes (Reifová et al., 2023). Consequently, we expect that genomic incompatibilities in hybrids will manifest early in haploid gametophytes as high mortality, whereas in diploid sporophytes, these barriers will more frequently emerge later, disrupting meiosis and life-cycle progression. Our aim here is to determine the relative impact of these barriers and whether meiotic failure truly represents an impermeable barrier to gene flow, using the model brown algae *Ectocarpus* (Peters et al., 2004).

**Figure 1.**
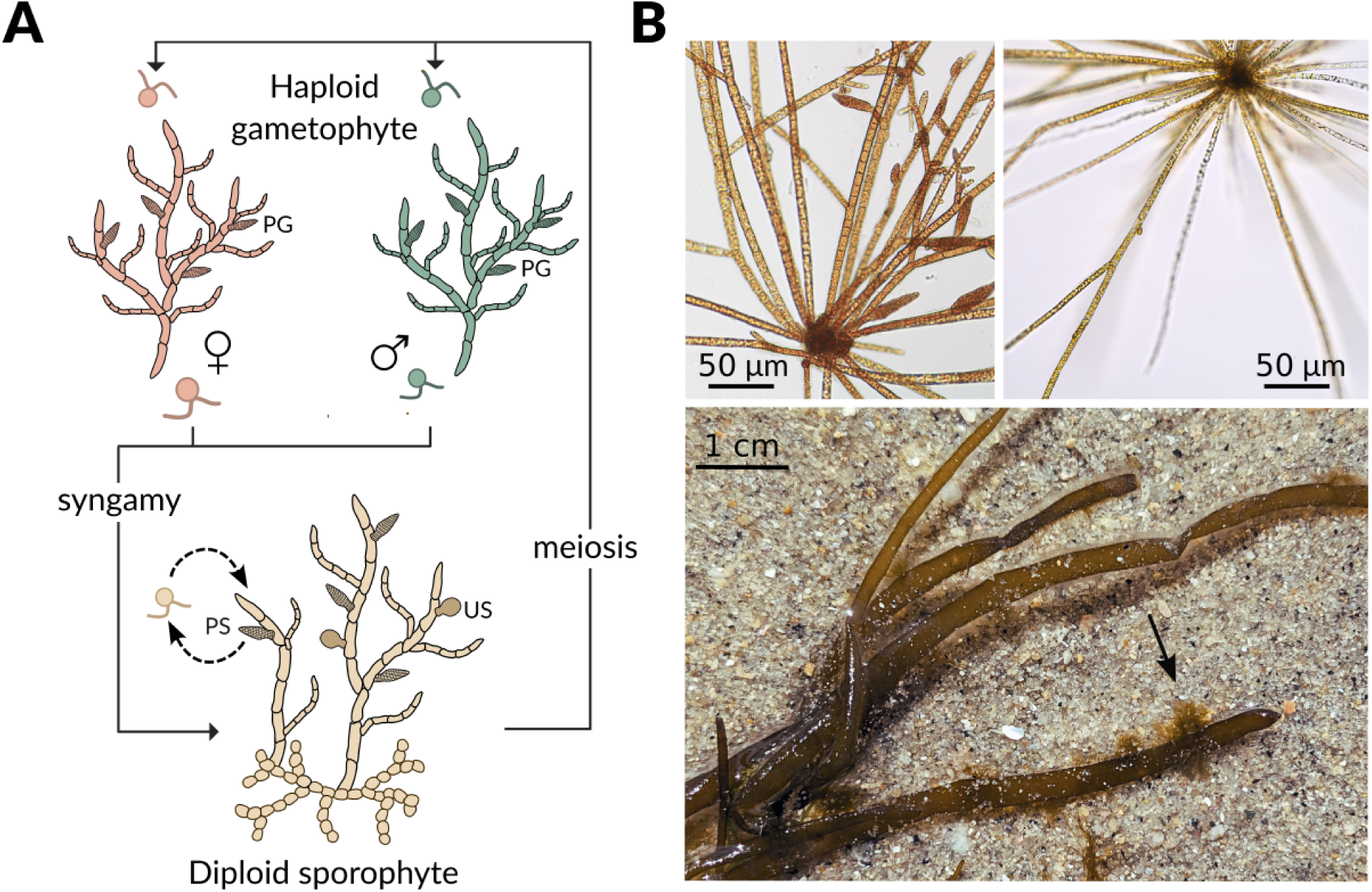
A) The *Ectocarpus* life cycle. The diploid sporophyte produces meio-spores through meiosis inside the unilocular sporangium (US). The released meio-spores develop as either male or female haploid gametophytes, which produce gametes in plurilocular gametangia (PG). Zygotes are formed by fusion of male or female gametes and develop as diploid sporophytes to complete the sexual cycle. In addition, diploid sporophytes produce mito-spores in plurilocular sporangia (PS), which germinate to produce a new diploid sporophyte generation (dashed line). B) Gametophytes of *Ectocarpus siliculosus* (top left) and *Ectocarpus crouaniorum* (top right). *Ectocarpus crouaniorum* (indicated by arrow) growing on *Scytosiphon* sp. (bottom).

*Ectocarpus* is a cosmopolitan filamentous seaweed, characterized by a haploid-diploid life cycle. While *Ectocarpus* was historically considered a pair of widespread species (Müller & Eichenberger, 1995), it is now recognized as a complex of at least 15 cryptic species with a history of complex divergence, strong reproductive barriers and occasional hybridization (Montecinos, Couceiro, et al., 2017). In the absence of obvious phenotypic differences, these species provide an excellent framework to dissect the genetic and ecological mechanisms that drive divergence along a speciation continuum, without confounding effects of morphological adaptation. This study focuses on species representing the two major *Ectocarpus* clades: *E. siliculosus* and *E. crouaniorum* (Fig. 1B) (Montecinos, Couceiro, et al., 2017; Peters et al., 2010). The two species exhibit distinct but overlapping distributions along the intertidal gradient, which could act as a partial ecological barrier. *Ectocarpus crouaniorum* is primarily found in high intertidal pools and frequently grows as an epiphyte on *Scytosiphon lomentaria* (Fig. 1B) (Couceiro et al., 2015; Peters et al., 2010). *Ectocarpus siliculosus* is mostly present in mid-intertidal to subtidal zones and has a broader range of host species (Couceiro et al., 2015; Peters et al., 2010). A large-scale study of European populations using microsatellite markers estimated that the two species hybridize at a level of 8.7%, with hybrids occurring only in sympatric zones and being almost exclusively diploid (Montecinos, Guillemin, et al., 2017). In controlled crosses, most interspecific zygotes aborted within 3-4 days and rare sporophytes that were raised to maturity were sterile and unable to complete meiosis (Peters et al. 2010). However, previous investigations of this hybrid zone relied on a limited number of microsatellite loci, which lack the resolution to fully characterize the genomic landscape of introgression.

Here we use high-resolution genomic data generated through linked-read sequencing (haplotype tagging or haplotagging) to overcome these limitations (Meier et al., 2021). This approach allows for physical phasing into large haplotype blocks, which is essential for visualizing the genomic ancestry of hybrids. As haplotagging has so far been applied to only a limited number of organisms, such as *Heliconius* spp. (Meier et al., 2021), this study represents yet another empirical application of the technique to a non-model system, such as populations of marine macroalgae. Moreover, by using whole genome resequencing data, we can move beyond simple hybrid identification to reconstruct a detailed population structure and infer the demographic history of the studied species. We replicated the previous European study by adding Chilean populations to test the universality of the reproductive barriers and determine if they are intrinsic or dependent on local geographic history and environmental factors. Our main aim is to shed light on how genomic differentiation, hybridization, and life-cycle complexity interact to shape speciation in marine macroalgae.

## 2. MATERIAL & METHODS

### Dataset definition

Our original dataset comprised 188 samples (including one additional technical replicate for 12 individuals) originating from Europe (n = 85) and Chile (n = 91) across different geographic regions (Table S1). Initial lineage assignment was based on nuclear ITS1 barcode polymorphisms and previous microsatellite data (Montecinos, Guillemin, et al., 2017). Using this approach, we identified 101 individuals as *E. crouaniorum* (Ecro) and 42 as *E. siliculosus* (Esil), with 14 individuals representing putative hybrids (Table S1). Ploidy and sex of the individuals were determined by PCR using male and female sex-specific markers (Table S2). Individuals carrying both sex markers were identified as diploid sporophytes, whereas individuals carrying only one marker were classified as male or female haploid gametophytes.

### Haplotagging library preparation and sequencing

High molecular weight DNA required for the haplotagging protocol was extracted following the method of Russo et al., (2022). The integrity of DNA was verified using a Femto Pulse, and DNA concentration was quantified using Qubit before dilution to 0.15 ng/uL (10 mM Tris, supplemented with 5 uL of 10mg/ml RNAse-A per 50 ml). Each DNA molecule was next labeled with a unique combination of four segmental barcodes (“A1-96”, “B1-96”, “C1-96”, “D1-96”) during the tagmentation reaction. Tagmentation was conducted using 2 µL (0.3 ng) of gDNA and 1.25 µL of haplotagging beads per sample, following the haplotagging protocol recommendations (Kucka & Chan, 2024). The 96-well plate of haplotagging beads was already assembled as described by Meier et al.,(2021). This combinatorial system enables the tagging of DNA molecules with up to 885,000 unique barcodes per sample of a 96-well plate. To increase multiplexing capacity, the 5th (plate) barcode (i7 index) was added during PCR, allowing us to include the whole dataset (188 samples) in one haplotagging library.

The tagmentation reaction was performed at 55°C for 10 min, then stopped with a stripping buffer (30mM NaCl, 10mM Tris, pH=8, 0.6% SDS). Un-integrated barcodes were removed from the bead surface through exonuclease 1 treatment. Tagmented DNA was next 5.3x subsampled before the PCR amplification. Subsampling step allowed us to preserve haplotagging bead diversity and target the optimal amount of DNA into the library (0.056 ng per sample).

PCR products were assessed on an agarose gel prior to size selection of the library using AMPure magnetic beads, targeting a range of 300-800 bp. The library was ultimately sequenced in-house on a P3 flow cell on a NextSeq 2000, generating 360 Gbp of data. Raw BCL files were converted to FASTQ format using bcl2fastq software, and reads were demultiplexed based on molecule-specific BX barcodes to assign linked reads to individual molecules and samples.

### Read mapping, SNPs calling and genotype imputation

Sequencing adapters were removed using cutadapt, and the reads were mapped to the chromosomal-level genome assembly of *Ectocarpus sp.7* (Cormier et al., 2017) using BWA (Li & Durbin, 2009). PCR duplicates were removed using Picard/markduplicate (https://broadinstitute.github.io/picard/). SNPs were called using samtools and bcftools (Danecek et al., 2021). Following quality control, only diploid individuals classified as *Ectocarpus* were retained. The diploid sporophytes were identified based on read coverage of the male and female sex-determining regions. For the 12 individuals sequenced in duplicate, we selected the replicate with better mapping statistics and lower genotype missing rate. After filtering, 166 individuals remained with an initial dataset containing 33.9 millions SNPs.

Genotype imputation was performed using STITCH (Turbek et al., 2021) by retaining exclusively biallelic SNPs with a mapping quality >20 and sequencing depth >10, reducing the initial VCF to 9,599,414 SNPs. To reduce computational time, each chromosome was divided into four intervals, and STITCH was run independently for each interval using all bam files as input with the following parameters: K=30, nGen= 200, expRate=15, shuffle_bin_radius=500, readAware=TRUE. The parameter K, representing the number of ancestral haplotypes, was optimized by testing multiple values and evaluating both the distribution of the imputation info score (IS) and the genotype concordance among the eight replicate pairs (Fig. S1). This benchmarking step allowed us to assess if the number of individuals and read coverage in the dataset were sufficient to correctly impute missing genotypes. The eight individuals with higher sequencing depth (5.8-8.3x) represented both species and potential hybrids, were next downsampled using Picard to 0.05X, 0.15X, 0.45X and 0.75X to mirror the lowest coverages in the dataset. We then used STITCH on a subset of chromosomes (1-4) with different parameters: K (5, 10, 20, 30 and 40), nGen (20 for K=10, 200 for all other values of K) to compare the genotype concordance, the calling rate (percentage of retrieved genotypes) and the distribution of the quality info score (IS) between original and downsampled datasets. To check for the genotype concordance and calling rate we focused on SNPs with IS > 0.4, AD > 5 or 10, and a quality ≥ 40.

After imputation, the average missing rate decreased from 43% to 25%, excluding SNPs with IS ≤ 0.5. After this step, we removed six individuals with >45% missing data. Final SNP filtering retained variants with genotype probability (GP) ≥ 0.92 and allowed ≤15% missingness, which resulted in a dataset of 160 individuals (Table S1) used for subsequent SNP-based and haplotype-based analyses.

### Species delineation using mitochondrial genomes

Using the unique segmental barcodes associated with linked reads, we reconstructed circularized mitogenomes for 154 out of the 160 individuals retained after mapping and filtering step using NovoPlasty v4.3 (Dierckxsens et al., 2017) and part of the COX1 gene (FP885846.2) as a seed. Circularized mitogenomes were aligned using Clustal Omega (Sievers & Higgins, 2018). A Neighbor-joining tree was constructed using MEGA V11.0.13 (Tamura et al., 2021), with *Ectocarpus sp7* (the reference genome used in this study) and *Pilayella littoralis* as an outgroup. Branch lengths were scaled proportionally using the 2-parameter Kimura method, systematically excluding ambiguous positions for each sequence pair. Mitochondrial phylogenies allowed confirmation of species identities. Using the same approach, we constructed a phylogenetic tree including individuals from the putative unknown species, individuals from our two species of interest as well as mitochondrial genomes from 12 cryptic species of *Ectocarpus* described in (Denoeud et al., 2024), and *Pilayella littoralis* as an outgroup (Fig. S2).

### Genetic structure

To minimize the effects of physical linkage among SNPs, we retained one variant every 100 bp, yielding 1.3 million SNPs for downstream analyses. Using this dataset, we inferred population structure using NGSadmix (MAF ≥ 0.01) implemented in ANGSD (Korneliussen et al., 2014), which took into account the genotype likelihood of each SNPs. To discern optimal clustering patterns within the dataset, we explored varying cluster numbers (K) ranging from 1 to 10, using 10 iterations per K. Genome-wide nucleotide diversity (π), genetic differentiation among lineages (Weir and Cockerham’s F_ST_), and the average number of nucleotide differences per site between populations (D_xy_) were estimated using Pixy (v1.2.7) (Korunes & Samuk, 2021) from a VCF file including invariant sites. Net nucleotide divergence between populations (D_a_) was calculated as D_xy_minus the mean within-population nucleotide diversity (π).

### Diagnostic SNP dataset and inference of hybrid class

To identify hybrid individuals, we first defined parental lineages based on ancestry estimates from NGSAdmix at K = 2 (>95% ancestry from one lineage) and species delimitation analyses based on mitogenomes (Fig. 2A,B). Following these criteria, 95 individuals of *E. crouaniorum* and 33 of *E. siliculosus* were identified as parental lineages. In addition, we defined first generation hybrids (putative F1-type and F1-like) by having approximately 45%-55% ancestry from either parental species, and admixed *E. siliculosus* individuals showing admixture from *E. crouaniorum* (5-20%). The proportion of hybrids was assessed separately within each geographic region to detect local variation in hybridization rates. We also examined the placement of mitogenomes from putative hybrids in the mitochondrial tree (Fig. 2B) to infer the maternal parent, as mitochondria are maternally inherited. This approach allowed us to assess the extent of genetic mixing between the two species and to evaluate the asymmetry of hybridization.

**Figure 2.**
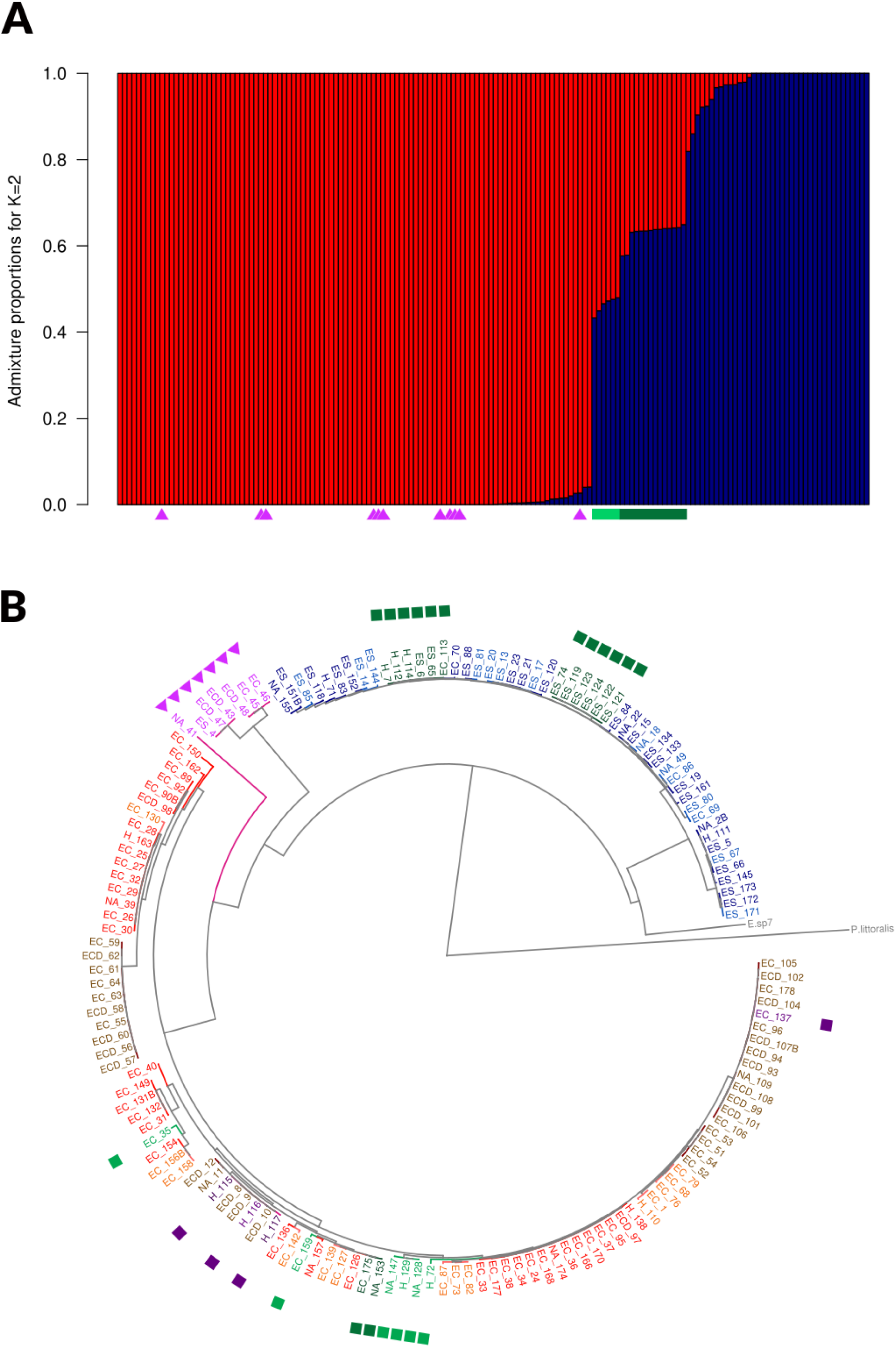
Cryptic diversity and hybridization within the *Ectocarpus* species complex. (A) Nuclear genetic structure based on NGSadmix analysis (K=2, n=160) showing the two parental species (*E. crouaniorum* in red and *E. siliculosus* in blue). The magenta cluster indicates an undescribed lineage (marked by triangles). Individuals with mixed ancestry indicate hybrids between *E.crouaniorum*-*E.siliculosus* (green). (B) Neighbor-Joining tree of mitochondrial genomes. The tree depicts the relationships among 154 out of the 160 individuals from this study, with *Ectocarpus sp7* (the reference genome species) and *Pilayella littoralis* as outgroups. Individual names are coloured according to genomic cluster: *E. siliculosus* (shades of blue), *E. crouaniorum* (shades of red), putative undescribed species (magenta, marked by triangles), *Esil* × *Ecro* hybrids (green, marked by circles), and hybrids between the unknown species and *E. crouaniorum* (purple, marked by circles).

To further describe hybridization patterns, we identified species-specific diagnostic SNPs for lineage pairs exhibiting admixture (Ecro1 vs Esil in Chile and Ecro3 vs. Esil in Europe, see Fig. 3). Diagnostic SNPs were identified by selecting loci with complete differentiation between parental groups (Weir and Cockerham’s F_ST_= 1) using VCFtools (Danecek et al., 2011). SNPs were further filtered to retain only high-quality genotypes (GP ≥ 0.96 instead of GP ≥ 0.92), ≤15% missing genotypes across individuals and one SNP every 1 kb. This resulted in a set of species-specific 5,871 SNPs (average distance = 32.9 kb) for the Chilean and 2,479 SNPs (average distance = 76.9 kb) for the European populations. Individuals with more than 40% missing data were excluded (9 from Chile, 11 from Europe).

**Figure 3.**
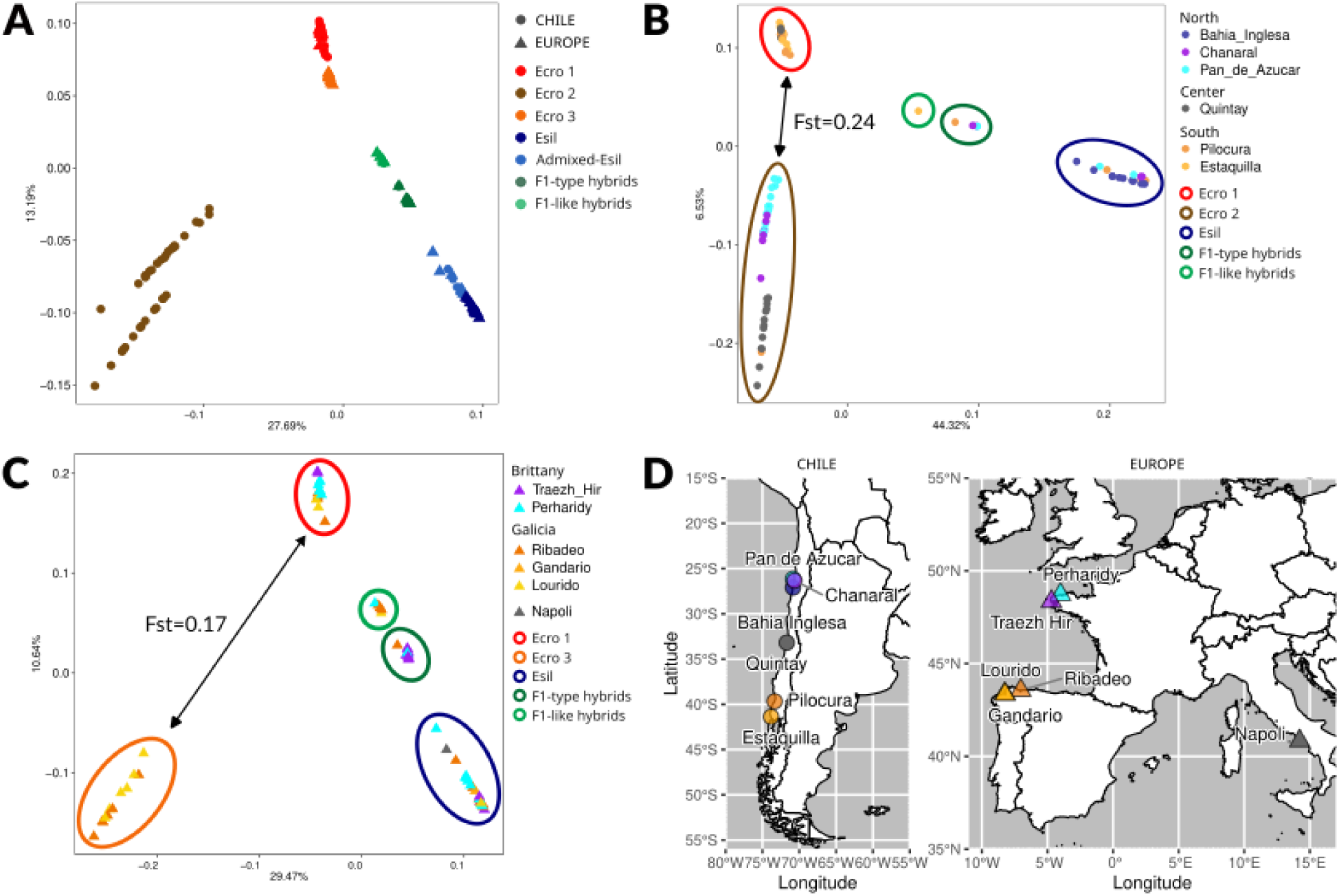
Population genomic structure and geographic distribution of *E. crouaniorum* and *E. siliculosus*. A) Global Principal Component Analysis (PCA) based on genotype likelihoods (n=149) showing lineage differentiation between the two *Ectocarpus* species and after removing the unknown species. *E. crouaniorum* forms three distinct clusters (Ecro1, Ecro2, and Ecro3), and putative hybrids are highlighted in green. Shapes denote geographic origin: the triangle displays European individuals and circles Chilean ones. B) Regional PCA of Chilean populations, illustrating the separation of the two Ecro clusters (Ecro1 and Ecro2). C) Regional PCA of European populations, showing the distribution of the Ecro clusters (Ecro1 and Ecro3). D) Map of sampling locations, displaying the collection sites across Europe and Chile included in this study. Points are color-coded by sampling locations.

The diagnostic SNP dataset was used to estimate inter-specific heterozygosity as well as a hybrid index defined as the admixture proportions derived from NGSAdmix at K = 2. Triangle plots were generated using inter-specific heterozygosity and the hybrid index (proportion of parental alleles), and observed genotypes were compared to those simulated under various hybridization scenarios. Simulations included F1, F2, F3, backcrosses (BC) with *Ectocarpus siliculosus* (Esil) or *Ectocarpus crouaniorum* (Ecro), spanning from one generation to four generations of backcrossing, “F1-BC type” (first and second generation of backcrosses with F1), “double backcross type” (first generation of backcrosses with the opposite pure species once, twice or third generation of backcrosses with the opposite pure species once or F1-BC type with the opposite species than the one for the backcross) (Fig. S3). For each hybridization scenario, we simulated 10 individuals. Simulations and their incorporation in the dataset, were conducted using the packages *vcfR* (Knaus & Grünwald, 2017), *adegenet* (Jombart & Ahmed, 2011) and *radiator* (Gosselin et al., 2020) through the ‘hybridize’ function from the diagnostic SNP VCF. Hybrid classifications were further validated using NewHybrids (Anderson & Thompson, 2002), which estimates posterior probabilities for parental, F1, F2, and backcross classes (Table S3) and their ancestry profile was compared to those obtained from haplotype information through the fineSTRUCTURE matrix (see ‘Haplotype-based inference of admixture’ below).

### Haplotype phasing and haplotype-based inference of admixture

Haplotype phasing was performed using the 10X Genomics Linked-Reads pipeline available with HapCUT2 (Edge et al., 2017). Phasing was conducted at the individual level over 500 kb intervals and involved several steps. First, BAM files were converted into fragment files using the extractHAIRS utility with the --10x option. Next, unlinked fragments were combined into barcoded molecules using LinkFragments.py with BX tag information, setting a maximum spacing of 50 kb between reads with the same barcode to consider them part of the same molecule. Finally, heterozygous positions were phased using HapCUT2 with the parameters --nf 1, --threshold 30, --error_analysis_mode 1, and --call_homozygous 1. Rather than performing subsequent statistical phasing steps, which can increase phasing errors, we retained the physical phasing strategy and then identified haplotype blocks consistently shared among individuals. This process involved first removing individuals with low phasing rates, then screening VCF files for positions with consistent PS tags, and finally detecting shared phased blocks across individuals. These consistently phased positions were then used to generate BED files for downstream analyses, resulting in the identification of 284 blocks (average block length 4.1 kb) that were phased consistently, with no missing data, across 57 individuals. Additional phased datasets were created for different lineage pairs to provide finer resolution of haplotype-based admixture (Table S4). Haplotype-based inference of admixture was performed using fineSTRUCTURE (Lawson et al., 2012). This process first involved using ChromoPainter to infer haplotype ancestry, which was then summarized in a co-ancestry matrix to quantify haplotype sharing. fineSTRUCTURE was run on five datasets: one previously defined dataset of 284 blocks across 57 individuals, and four additional datasets corresponding to individual lineage pairs (see Table S4).

### Demographic inference (DILS)

Scenarios of speciation were evaluated using the Demographic Inferences with Linked Selection (DILS) pipeline (Csilléry et al., 2012; Fraïsse et al., 2021; Pudlo et al., 2016). DILS is an ABC-based approach that compares summary statistics of genomic variation to simulations under different demographic scenarios. Specifically, in the two-population mode, DILS uses a hierarchical model-selection procedure to distinguish between models of current isolation (SI: strict isolation; AM: ancient migration) and models with ongoing migration (IM: isolation with migration; SC: secondary contact). We ran DILS for each pair of *E. siliculosus* and *E. crouaniorum* lineages (in Chile: Ecro1-Esil and Ecro2-Esil; in Europe: Ecro1-Esil and Ecro3-Esil) to first evaluate the extent of gene flow within each pair, and then to determine whether lineage divergence has occurred in parallel across geographic regions and lineages. F1-hybrids and admixed individuals detected previously were removed from each dataset. Since DILS currently handles a maximum of 1,000 loci, inferences were made by subsampling genomes into 2 kb segments, ensuring that segments were separated by at least 30 kb before randomly selecting 1,000 loci (167,781 SNPs in total). The same set of loci was consistently used across lineage pairs DILS analyses were run under a heterogeneous effective size model (hetero-Ne) employing the default priors listed in Table S5.

### Localization of barriers to gene flow

To identify genomic regions associated with reproductive barriers, we performed chromosome-level painting of individual genomes based on ancestry inference using the EM algorithm implemented in *diem* (Baird et al., 2023). This method infers allele ancestry directly from genotype data and does not require predefined reference individuals or candidate variants. At each variant position, genotypes are assigned as 0/0, 1/1, or 0/1, corresponding to homozygosity for either species (group of individuals without priors) or heterozygosity, respectively. To reduce noise caused by shared polymorphisms or low-confidence sites, we selected only the 40% of markers with the highest diagnostic index. The polarization analysis was conducted for the complete dataset of 149 individuals and also focusing specifically on the hybrid individuals to characterize the genomic distribution of introgressed regions.

## 3. RESULTS

### Hidden diversity within cryptic species of *Ectocarpus*

While this study initially aimed to investigate reproductive isolation between two closely related *Ectocarpus* species, our genomic analyses revealed a more complex pattern of cryptic diversity than initially assumed. Principal component analysis (PCA) and NGSadmix analyses based on unlinked SNPs for the initial dataset of 160 individuals identified two major genetic lineages corresponding to *E. siliculosus* (Esil, in blue) and *E. crouaniorum* (Ecro, in red), as well as a group of intermediate individuals (in green) which were later identified as hybrids between these two species (Fig. 2). Notably, we identified a cluster of distinct 11 individuals (10 in Europe, one in Chile) half of which showed admixture with *E. crouaniorum* (unknown species in magenta, Fig. 2A,B). Genetic structure analyses further confirmed the occurrence of this third unknown lineage, which explained 41.56% of the total variance along PC1 in Europe (Fig. S4A). This lineage showed strong differentiation from *E. siliculosus* (F_ST_ = 0.72) and moderate differentiation from *E. crouaniorum* (F_ST_ = 0.51). Comparison with the previously published *Ectocarpus* mitogenomes (Denoeud *et al*. 2024) suggests that this lineage likely represents an undescribed species that is phylogenetically closer to *E. crouaniorum* than to *E. siliculosus* (Fig. S2 and S4B).

The phylogenetic analysis of reconstructed, circularized mitogenomes were in congruence with the nuclear data and identified two major groups corresponding to our focal species independent of geographical origin, with the unknown species branching off the *E. crouaniorum* lineage (Fig. 2B). Putative hybrids between *E. crouaniorum* and *E. siliculosus* identified in the admixture plot (Fig. 2A), were placed in both mitochondrial clades (in green Fig. 2B). Furthermore, the five individuals showing nuclear admixture between the undescribed lineage and *E. crouaniorum* all possessed *E. crouaniorum*-type mitochondria, providing further evidence of ongoing hybridization between these two lineages (in purple, Fig. 2B). To ensure a clear delimitation of the hybrid zone between *E. siliculosus* and *E. crouaniorum* and to avoid confounding signals of introgression of the undescribed species with *E. crouaniorum*, individuals belonging to the unknown species (i.e., 11 individuals) were excluded from the subsequent analyses.

Next, we analyzed the population structure within *E. crouaniorum* and *E. siliculosus* (Fig. 3). PCA and NGSadmix analyses revealed that the *E. crouaniorum* lineage consists of three differentiated clusters (hereinafter Ecro1, Ecro2, and Ecro3) whereas *E. siliculosus* showed limited genetic variation, forming a single, cohesive lineage (Esil) in Chile and Europe (Fig. 3A). In Chile, *E. crouaniorum* formed two geographically differentiated clusters, separating northern (Ecro2) and southern (Ecro1) populations, with individuals from central Chile represented in both clusters (Fig. 3B,D). In Europe, *E. crouaniorum* also formed two distinct genetic clusters with geographic separation (Ecro1 and Ecro3), individuals from Brittany were restricted to Ecro3 while both, Ecro1 and Ecro3 clusters were observed in Galicia (Fig. 3C,D). In contrast, *E. siliculosus* individuals formed a single genetic group independent of geographic location.

Together, these analyses reveal a complex evolutionary landscape involving three distinct genetic clusters within *E. crouaniorum*, an additional unknown cryptic species, and evidence of ongoing hybridization.

### Geographic and lineage-specific patterns of hybridization

Genomic admixture and hybridization between *E. crouaniorum* and *E. siliculosus* varies between clusters and geographic regions. Signals of interspecific contact were more frequent in Europe (22 individuals showing admixture (> 5% ancestry proportion), 56% of all samples) than in Chile (12 individuals; 31% of all samples) (Fig. 4A,B). Within these admixed individuals, we identified both high-ancestry hybrids as well as individuals showing lower levels of admixture, although these involve different *E. crouaniorum* clusters in each region. In Europe, Ecro3 hybridizes with *E. siliculosus* while Ecro1 remains isolated. In contrast, in Chile, Ecro1 hybridizes with *E siliculosus* while Ecro2 does not (Fig. 3B,C; Fig. 4A,B). This pattern is partially explained by genetic divergence, where D_a_ and D_xy_ are higher between non-hybrydizing Ecro1 and *E. siliculosus* (D_a_= 0.029, D_xy_ = 0.037, F_ST_ =0.68) than between the hybridizing Ecro3 and *E. siliculosus* (D_a_= 0.025, D_xy_= 0.032, F_ST_ =0.71) in Europe, and between the non-hybridizing Ecro2 and *E. siliculosus* (D_a_=0.034, D_xy_ = 0.049, F_ST_ = 0.42) compared to hybridizing Ecro1 and *E. siliculosus* (D_a_=0.027, D_xy_ = 0.036, F_ST_ = 0.67) in Chile. Remarkably, the Ecro1 clusters from Chile and from Europe are nearly identical (F_ST_ = 0.01) and the same pattern is observed for Esil, with Chilean and European populations also showing minimal differentiation (F_ST_ = 0.01), suggesting that the barriers to gene flow likely depend on the environment or differences in gamete compatibility and fertilization dynamics rather than intrinsic genomic incompatibilities.

**Figure 4.**
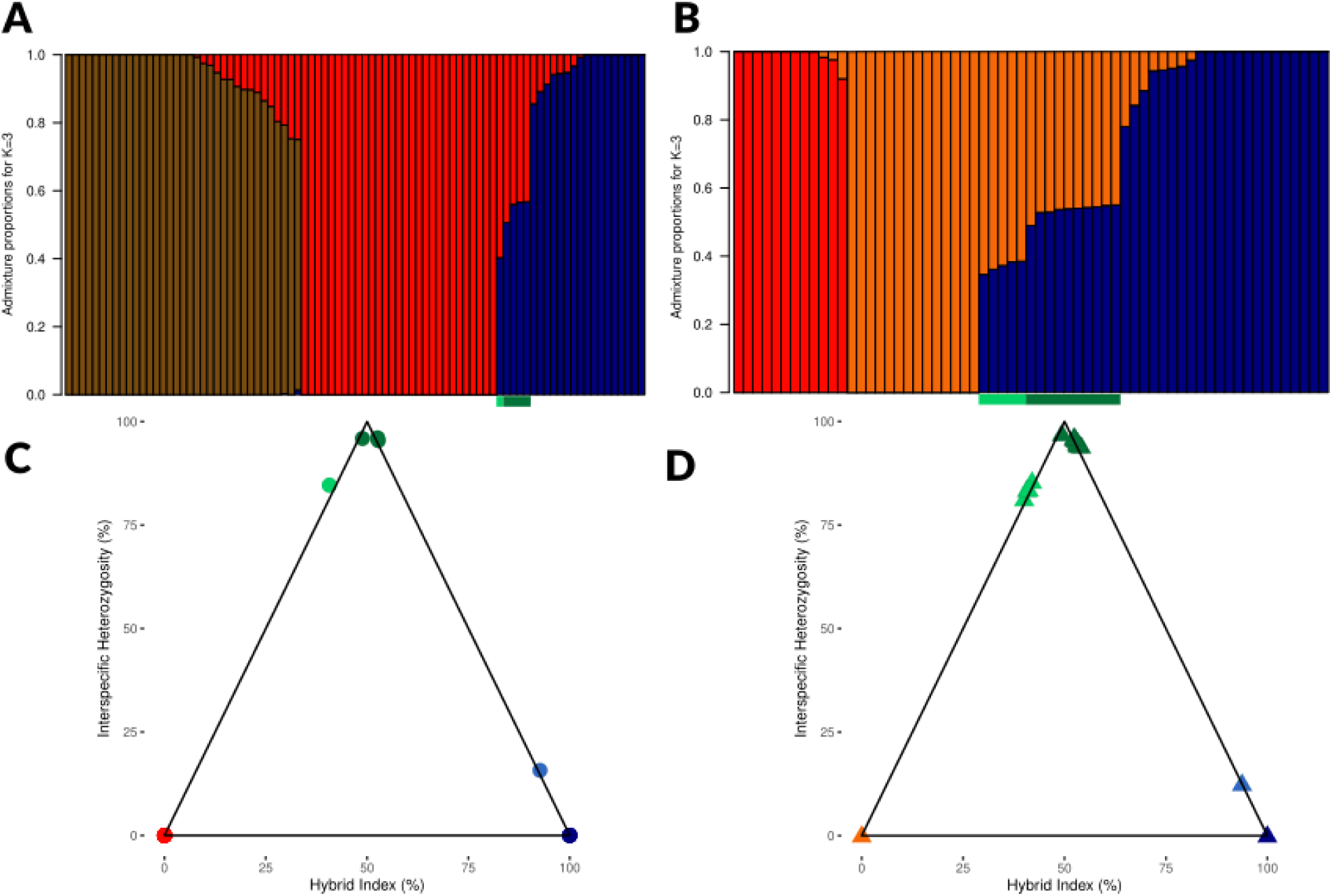
Genomic admixture and hybrid classification in European and Chilean *Ectocarpus* populations. A-B) Admixture results using K=3 for (A) 86 Chilean individuals and (B) 63 European individuals. Color code follows the results of NGSadmix and PCA per geographic region (Europe vs. Chile); *E. siliculosus* in blue, *E. crouaniorum* clusters (Ecro1: red; Ecro2: brown; Ecro3: orange), putative hybrids in green. C-D) Triangle plots of hybrid index vs. interspecific heterozygosity. The x-axis represents the Hybrid Index (proportion of *E. siliculosus* alleles), ranging from *E. crouaniorum* (0.0, left) to *E. siliculosus* (1.0, right). (C) 42 Chilean individuals analyzed using 5,871 species-specific SNPs between *Ecro1* and *Esil*. (D) 40 European individuals analyzed using 2,479 species-specific SNPs between Ecro3 and Esil. Points are color-coded by genomic category: pure *E. siliculosus* (blue), pure *E. crouaniorum* (red), F1-type hybrids (dark green), F1-like individuals with excess of *E. crouaniorum* alleles (light green), and admixed *E. siliculosus* individuals (light blue).

To further characterize the hybrids, we analyzed interspecific heterozygosity and hybrid indices, and performed simulations (Fig. S3) to categorize them into three distinct classes: 1) F1-type hybrids, 2) F1-like individuals with reduced heterozygosity and excess of *E. crouaniorum* alleles, 3) admixed *E. siliculosus* with introgressions from *E. crouaniorum*. Because hybrid classification using triangle plots required filtering for individuals with less than 40% missing data at species-diagnostic SNPs, the number of individuals that could be confidently assigned to hybrid classes was reduced compared with the initial admixture analysis (Fig. 4). However, the detected hybrids were fully consistent across the PCA, triangle plot, fineSTRUCTURE, and admixture analyses.

The most prevalent class in both Chile and Europe were the F1-hybrids (Fig. 4C,D). In Chile we identified three F1-type hybrids (Class 1), one F1-like individual with excess of *E. crouaniorum* alleles (Class 2), and one admixed *E. siliculosus* (Class 3) (Fig. 4C). In Europe, we identified nine F1-type hybrids (Class 1), four F1-like individuals with excess of *E. crouaniorum* alleles (Class 2), and one admixed *E. siliculosus* (Class 3) (Fig. 4D). Additional individuals showing some levels of admixture (<20%) were observed in the NGSadmix analyses (Fig. 4A,B), but these samples exhibited low sequencing coverage and high levels of missing data and therefore did not pass the filter threshold for the triangle plot analysis. Because hybrid index estimation relies on accurate genotype calls at species-diagnostic loci, missing data and genotype imputation can artificially move individuals toward intermediate positions in the triangle plot, which could lead to false inference of hybrid class. These individuals were therefore excluded from the hybrid classification. However, the presence of admixed individuals that were identified with high confidence (Fig. 4C,D) suggests that the meiotic barrier in F1 sporophytes is not absolute and that introgression between the species can occur.

Finally, the mitochondrial analysis (Fig. 2B) showed that while F1 hybrids predominantly possessed *E. siliculosus* mitochondria, seven individuals carried the *E. crouaniorum* haplotype, which was detected in both Europe and Chile. This suggests that hybridization is not strictly unidirectional, as the mitochondrial genome, which is uniparentally inherited from the mother, could be derived from either parental species.

### Demographic inferences of gene flow and genomic barriers

The extent of reproductive isolation was investigated using the Demographic Inferences with Linked Selection (DILS) pipeline (Fraïsse et al., 2021) among lineage pairs showing different potential of interspecific gene flow and distinct levels of genomic divergence. In Europe between hybridizing lineages (Ecro3-Esil) and non-hybridizing lineages (Ecro1-Esil) and in Chile between hybridizing lineages (Ecro1-Esil) and non-hybridizing lineages (Ecro2-Esil). DILS analyses supported scenarios of ongoing migration for all tested lineage pairs (posterior probability = 0.89-0.92, Table 1). The migration patterns were heterogeneous across loci with high posterior probabilities (0.69-0.86, Table S6), allowing to further classify them as barrier-associated loci and migration-associated loci (Table 1, Table S7).

**Table 1.** Demographic inference (DILS) and genomic isolation patterns.

| Pairwise comparison | Region | Hybrids detected | Model | Tsplit | $M1 \rightarrow 2$ | nBarrier 1→2 | $M2 \rightarrow 1$ | nBarrier 2→1 | %Barrier loci | % Migration loci |
| --- | --- | --- | --- | --- | --- | --- | --- | --- | --- | --- |
| Ecro3-Esil | Europe | Yes | IM | 70,052<br>[50,142-96,793] | 68<br>[42-92] | 225<br>[174-295] | 3.50<br>[199-5.10] | 225<br>[135-324] | 14.1 | 59.8 |
| Ecro1-Esil EU | Europe | No | SC | 218,708<br>[169,726-288,254] | 47<br>[34-64] | 312<br>[242-381] | 4.58<br>[2.63-6.74] | 202<br>[124-291] | 19.3 | 68.5 |
| Ecro1-Esil CH | Chile | Yes | IM | 289,883<br>[211,221-359,216] | 14<br>[8-20] | 401<br>[291-536] | 24<br>[16-33] | 141<br>[94-193] | 10.6 | 83.5 |
| Ecro2-Esil | Chile | No | IM | 738,530<br>[410,247-1,147,666] | 63<br>[47-81] | 296<br>[202-397] | 8<br>[5-11] | 0.53<br>[0-1] | 23.4 | 58.1 |

As expected, divergence time was positively correlated with the number of barrier loci between the interacting lineages (Table 1, Fig. S5). In Europe, the Ecro3-Esil pair represented the most recent split, with an estimated divergence time of ∼70,000 generations and a moderate proportion of barrier loci (14%) relative to migration loci (84%) (Table 1). This pair also exhibited the lowest level of genetic divergence (D_a_= 0.025, D_xy_ = 0.032) and showed the highest hybridization potential (Fig. 4B,D). In contrast, the non-hybridizing Ecro2-Esil pair, the oldest among the comparisons and estimated to have diverged approximately 739,000 generations ago, showed nearly double the proportion of barrier loci (23%) relative to migration loci (58%) (Table 1). These results illustrate the gradual accumulation of genomic barriers over hundreds of thousands of generations.

However, genetic distance alone does not fully explain reproductive success, as evidenced by the paradox of Ecro1-Esil interaction, which results in hybridization in Chile, but not in Europe. DILS analyses revealed that these lineages possess distinct evolutionary histories across regions. In Chile, an isolation-with-migration (IM) model was supported, displaying the lowest proportion of barrier (11%), relative to migration loci (84%) (Table 1). In contrast, in Europe, the secondary contact (SC) model was preferred, associated with the highest divergence (D_a_= 0.029, D_xy_ = 0.037) alongside almost double the amount of barrier loci (19%). This suggests that the period of allopatry between Ecro1 and Esil in Europe may have strengthened the barriers more effectively than the ongoing migration in Chile. Consequently, local demographic history, rather than intrinsic genetic distance alone, is the determinant of the successful hybridization in *Ectocarpus*.

Finally, DILS revealed that gene flow is highly asymmetric. In all comparisons except Ecro1-Esil in Chile, migration was heavily biased from *E. crouaniorum* into *E. siliculosus* (M1→2 up to 20-fold higher than M2→1) (Table 1). This directional bias is consistent with the genomic ancestry patterns observed in the admixture plots (Fig. 4 A,B). The only exception was the Ecro1-Esil in Chile where the migration was more balanced (M1→2=14; M2→1=24). These results suggest that overall *E. crouaniorum* maintains its genomic integrity more strongly than *E. siliculosus.* Since the number of loci was similar in both directions (Table 1), the difference in effective migration rates could be driven by ecological factors rather than solely intrinsic genomic incompatibilities. In addition, we observed a substantial amount of migration loci even in non-hybridizing species (58% in Ecro2-Esil in Chile and 68% in Ecro1-Esil in Europe). This pattern suggests that the absence of F1 hybrids in our dataset does not imply strict reproductive isolation and may instead reflect ancient introgression, rare hybridization events, or indirect gene flow. Taken together, these results show that reproductive isolation in *Ectocarpus* is a complex interplay between genetic divergence, demographic history and geographic context.

### The genomic landscape of hybridization

Admixture analyses based on both haplotype sharing and unlinked SNPs produced highly consistent results, revealing several clusters and sub-clusters within *E. crouaniorum* and strong differentiation from *E. siliculosus* (Fig. 5A). *E. crouaniorum* cluster 2 (Ecro2) is strongly differentiated from the two other *E. crouaniorum* clusters and highly divergent from *E. siliculosus*. This is in line with the oldest divergence time inferred by DILS (∼738,000 generations) (Table 1). Analyses of haplotype sharing further revealed internal genetic sub-structure consisting of two major sub-clusters within Ecro2 (Fig. 5A). In contrast, *Ecro1* from both Chile and Europe formed a single, cohesive cluster. Consistent with the higher number of hybrids observed in Europe, where Ecro3 hybridizes with *E. siliculosus*, Ecro3 showed significantly greater haplotype sharing with Class 1 (F1-type) and Class 2 (F1-like) hybrids than any other lineage.

**Figure 5.**
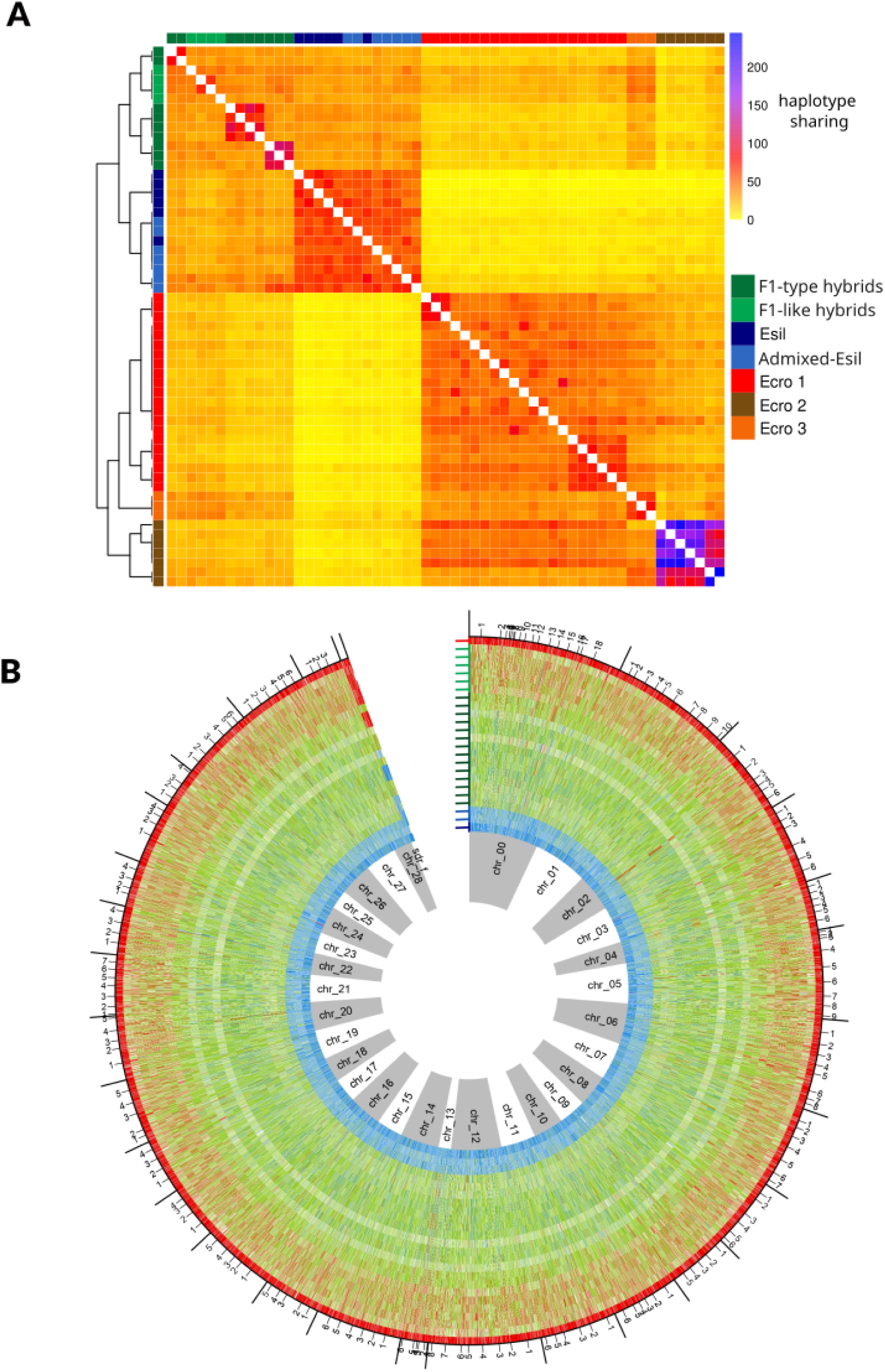
A) Haplotype-based inference of admixture, with the co-ancestry matrix and the inferred tree showing relationships between individuals. Rows represent recipients and columns represent donors. Haplotype sharing was first inferred with ChromoPainter using 284 phased blocks shared across a subset of 57 individuals. Colour intensity indicates the amount of haplotype sharing (copied chunks) inferred by ChromoPainter. The resulting co-ancestry matrix was then analysed with fineSTRUCTURE to cluster individuals based on their haplotype copying profiles. B) Circular representation of the location of barriers and the polarity of diagnostic markers for one parental species, admixed and hybrid samples (n=24) across chromosomes. The inner ring shows the location of each chromosome in alternating grey and white. Moving outwards, the next ring indicates one Esil parental species (in blue), then the two admixed Esil (light blue), the 14 F1-like (dark green), the six F1-type hybrids (light green), and one Ecro parental species (in red). Each ring shows the genotypes of each sample at each diagnostic marker (p>0.6).

To characterize the physical distribution of ancestry, we used the Diagnostic Index Expectation Maximization method (DIEM) (Baird et al., 2023). This analysis confirmed that Class 1 F1 hybrids maintained near-perfect heterozygosity across all diagnostic SNPs (Fig. 5B, Fig. S6). As expected, Class 2 individuals, with F1-like genomic architecture but an excess of *E. crouaniorum* alleles, showed slightly higher levels of haplotype sharing with *E. crouaniorum* relative to F1 of Class 1. In the admixed *E. siliculosus* individuals (Class 3), *E. crouaniorum* introgression was restricted to small segments scattered stochastically across the genome (Fig. 5B). We found no evidence of conserved introgression tracts among different individuals, which suggests that no specific genomic regions are preferentially favored by selection.

To summarize, we found that ancestry segments were inconsistently distributed across genomes of admixed individuals, representing rare, fragmented recombination events that are likely being purged or diluted over time.

## 4. DISCUSSION

### Genomic resolution of hybridization in *Ectocarpus*

The application of linked-read whole genome sequencing (haplotagging) greatly increased genomic resolution and revealed previously unrecognized levels of complexity within *Ectocarpus* hybrid zones. While previous studies relied on a limited set of nine microsatellite loci and single-gene barcodes (Montecinos, Guillemin, et al., 2017), our approach generated a massive increase in marker resolution, identifying over 1.3 million high-quality, unlinked SNPs across the genome. In addition, the use of linked sequence information enabled us to phase haplotype blocks and perform high-resolution ancestry painting to characterize introgression patterns genome-wide.

In particular, we detected six times more hybrids than previously reported (Montecinos, Guillemin, et al., 2017) and uncovered not only the presence of F1 hybrid sporophytes between *E. crouaniorum* and *E. siliculosus* but also *E. siliculosus* individuals carrying introgressed *E. crouaniorum* genomic segments at lower frequencies. Sixteen of these individuals showing clear signatures of admixture had previously been assigned to parental species based on microsatellite data (Montecinos, Guillemin, et al., 2017), underscoring the limited power of such markers to detect low levels of introgression. As shown here, introgressed tracts between the two *Ectocarpus* species are typically small and stochastically distributed across the genome, making them largely undetectable with sparse microsatellite loci.

One limitation of our study, however, is the use of a heterospecific reference genome (*Ectocarpus sp. 7*) (Cock et al., 2010; Cormier et al., 2017), as continuous genome assemblies for *E. crouaniorum* and *E. siliculosus* are not yet available. Reference bias may affect population genomic analyses due to sequence divergence, structural differences, and mapping inefficiencies between the focal and reference species (Prasad et al., 2022). Reference bias generally increases with phylogenetic distance and may inflate heterozygosity due to misalignments being called as variants (Akopyan et al., 2025; Prasad et al., 2022). In our case, the divergence between both focal species and *Ectocarpus sp.7* remains modest in the context of *Ectocarpus* phylogeny, which resulted in comparable read mapping efficiency for *E. siliculosus* and *E. crouaniorum* (55.5% and 49.3% respectively), and after quality filtering we recovered millions of SNPs that were uniformly distributed across the genome (Fig. S7).

Only two chromosomes showed significantly reduced coverage, one of which (chr13) corresponds to the sex chromosome across the brown algae (Ahmed et al., 2014; Barrera-Redondo et al., 2025), and which has previously been identified as a hotspot of divergence in *Ectocarpus* (Denoeud et al., 2024). While sex chromosomes are widely implicated in the evolution of reproductive incompatibilities (Coyne & Orr, 1989; Haldane, 1922; Payseur et al., 2018), the specific role of haploid UV sex-determination systems in speciation remains largely unexplored, which presents a compelling avenue for future research once species-specific assemblies are completed.

Because analyses of structural variants and large chromosomal rearrangements may be biased by the choice of reference (Maurstad et al., 2025), these patterns of genome evolution were not investigated here. While structural variants between *E. crouaniorum* and *E. siliculosus* could potentially contribute to reproductive incompatibilities, we focused our analyses on population structure, genetic differentiation, and demographic history. These analyses are robust to moderate reference divergence and ensure that the patterns of admixture and lineage differentiation reflect true biological signals.

### Cryptic species diversity and population structure

The increase in marker resolution allowed us to move beyond simple hybrid detection, revealing previously undescribed genetic structure in *Ectocarpus crouaniorum* and evidence of further cryptic diversity within the genus *Ectocarpus*. These genetic differences are not reflected in obvious morphological traits, a pattern which is increasingly observed in many macroalgal taxa as more genomic data becomes available (Hoshino et al., 2018; Knoop et al., 2024; Payo et al., 2013). Such hidden diversity has important implications for studies of gene flow and hybridization, as cryptic species may differ in their reproductive compatibility and demographic histories despite appearing morphologically identical (Christmas et al., 2021; Hoshino et al., 2021). For instance, we found a third unknown species of *Ectocarpus* in Europe that appears to hybridize with *E. crouaniorum*. The discovery of cryptic species that actively hybridize suggests that reticulate evolution in brown algae may be more pervasive than currently appreciated, and that unrecognized species boundaries can confound inferences of introgression and demographic history.

Recognizing cryptic diversity is also critical when studying adaptation and functional traits, as distinct lineages may differ in physiology, ecological tolerances, or metabolite production (Muangmai et al., 2015; Nygren, 2014; Scriven et al., 2016); reviewed in (Bickford et al., 2007)). For example, many cryptic macroalgae have been found to exhibit different physiological responses to wave exposure, depth, herbivory, or varying salinity and temperature levels (Martin et al., 2024; Muangmai et al., 2015; Puk et al., 2020). Our results show that *E. crouaniorum* comprises three genetically distinct clusters, whereas *E. siliculosus* appears to form a single, globally distributed lineage. This divergence may be driven by their contrasting ecological niches. *E. crouaniorum* typically inhabits the upper intertidal zone, where organisms experience strong environmental fluctuations, including heat stress, salinity variation, and wave exposure (Couceiro et al., 2015; Peters et al., 2010). Such heterogeneous and stressful environments may promote local adaptation and population divergence. In contrast, *E. siliculosus*, which generally occurs lower in the intertidal zone, may experience more stable environmental conditions, potentially reducing opportunities for ecological differentiation (Peters et al., 2010). Furthermore, populations located in the upper intertidal zone experience greater temporal and spatial isolation due to tidal cycles than those in lower shore habitats (Engel et al., 2004; Krueger-Hadfield et al., 2013). As a result, lower-shore populations tend to exchange more genes, whereas upper-shore populations are more genetically isolated. For example, in the red alga *G. gracilis*, F_ST_ values are generally low among tide pools in the lower intertidal zone, and high among those in the upper intertidal zone (Engel et al., 2004). Such reduced gene flow in upper intertidal habitats may promote local adaptation and, ultimately, facilitate ecological divergence and speciation. These observations fit in the framework of ecological speciation, where divergent selection in heterogeneous environments drives reproductive isolation (Shafer & Wolf, 2013). Such patterns have been documented in other intertidal organisms, such as *Littorina* snails, where environmental gradients across the shore drive rapid diversification (Johannesson et al., 2024; Stankowski et al., 2020).

### Replicated hybrid zones

To test whether ecological divergence consistently contributes to lineage isolation in *Ectocarpus,* we investigated barriers to gene flow across multiple geographic hybrid zones. Replicated hybrid zones represent a paradigm shift in speciation research, moving us from interpreting single, case-specific contact zones to testing how repeatable and predictable hybridization outcomes are across space and time (Westram et al., 2021). Studying multiple, independent contact zones between the same species, as in our *Ectocarpus* system, allows us to distinguish stochastic variation from deterministic responses to similar ecological and demographic conditions, thereby strengthening inferences about which barriers to gene flow are context dependent versus intrinsic (Castaño et al., 2026; Łabiszak et al., 2025; Pal et al., 2025).

In both Europe and Chile, *E. crouaniorum* and *E. siliculosus* form hybrid zones dominated by F1-type hybrids with evidence of asymmetric gene flow from *E. crouaniorum* into *E. siliculosus*, suggesting that broad aspects of hybrid zone structure and directional introgression can be repeatable, analogous to recurrent asymmetric introgression described in pines, birds and plants (Castaño et al., 2026; Łabiszak et al., 2025; Ocampo et al., 2023; Pal et al., 2025) or even red algae (Reynes et al., 2026). When similar patterns of admixture and biased migration occur repeatedly across regions, they point to predictable features of the speciation process under comparable ecological settings (Castaño et al., 2026; Łabiszak et al., 2025; Ocampo et al., 2023). At the same time, we showed that parallelism in broad patterns of hybridization (Ecro1-Esil in Chile, Ecro3-Esil in Europe) can coexist with non-parallel, context-dependent genetic outcomes (Ecro1-Esil in Europe and Chile). In Chile, *E. crouaniorum* cluster 1 (Ecro1) hybridizes with *E. siliculosus*, while in Europe, Ecro1 remains strictly isolated from *E. siliculosu*s, despite Chilean and European Ecro1 populations being nearly identical (F_ST_=0.01). Notably, demographic inference supports different histories in the two regions, with secondary contact being predicted in Europe and isolation with migration being the best scenario in Chile. These contrasting outcomes of species interaction suggest that ecological or demographic context, rather than intrinsic genomic incompatibility alone, determine whether a given lineage participates in hybridization. In particular, the period of allopatry inferred for European populations may have reinforced reproductive barriers, as observed in other systems (Augustijnen & Lucek, 2024; Deshmukh et al., 2025; Muto & Kai, 2023). Nevertheless, we also infer substantial migration between lineage pairs in which no current F1 hybrids are observed (Ecro2-Esil in Chile, Ecro1-Esil in Europe), suggesting that the absence of visible hybrids in a given transect does not necessarily imply strict reproductive isolation but may instead reflect rare, ancient, or episodic gene flow. Alternatively, gene flow between these species pairs may have ceased only recently, in which case DILS may lack the resolution to detect this change. We note that the distinction between "gene flow" and "secondary contact" models may partly reflect differences in the timing or duration of contact rather than fundamentally different modes of speciation, with hybrid frequency expected to be higher in the recent zone, as found in oaks (Liao et al., 2019).

This decoupling between short-term hybrid presence and long-term gene flow has been reported in other hybrid zones, where similar phenotypes or hybrid zone structures arise from distinct genetic architectures and evolutionary histories (Castaño et al., 2026; Ocampo et al., 2023; Pal et al., 2025). In these systems, hybridization outcomes are shaped by individual characters yet they converge on predictable features such as directionality of introgression and the repeated involvement of specific trait loci. Our results in *Ectocarpus* highlight how a comparative, multi-replicate framework reframes speciation as a balance between predictability and contingency. Evolution is predictable at the level of hybrid zone formation and asymmetry of introgression, but remains non-parallel at the level of which lineages hybridize and how the genetic architecture of reproductive isolation is assembled across different geographic and environmental settings.

### Genomic architecture of isolation

To further infer direction, localization and extent of gene flow, we examined the genomic architecture of reproductive isolation between the two species. We found that introgression blocks between *E. crouaniorum* and *E. siliculosus* were homogeneously distributed across the genome. This pattern is consistent with a polygenic barrier, where reproductive isolation is maintained by numerous loci of small effects (Kautt et al., 2020). Such an architecture was originally proposed as a null model of reproductive isolation (Barton & Charlesworth, 1984) and contrasts with the "speciation islands" model, which expects divergence to be concentrated in a few large-effect regions (Malinsky et al., 2015; Nguyen et al., 2024). Evidence of polygenic barriers to gene flow has recently been described for many taxa, including fish (Haenel et al., 2021), butterflies (Ebdon et al., 2025; S. H. Martin et al., 2019), or fruit fly (Morán & Fontdevila, 2014), suggesting this pattern may be more prevalent in speciation than previously thought (Kautt et al., 2020).

Polygenic barriers in *Ectocarpus* are supported by local ancestry inference, which revealed small, stochastically scattered, introgressed *E. crouaniorum* segments in admixed *E. siliculosus* individuals. The lack of conserved introgression tracts among different individuals suggests that no specific genomic regions are preferentially favored by selection, but rather that recombination fragments these segments while selection gradually purges them. However, the striking asymmetry of gene flow, strongly biased from *E. crouaniorum* into *E. siliculosus*, suggests that gene flow in the opposite direction is strongly selected against.

This directional bias may be explained by several non-mutually exclusive factors. Firstly, ecological differences between the species may play a major role. Although both are epiphytic, *E. crouaniorum* appears to be more host-specific, whereas *E. siliculosus* is a generalist (Couceiro et al., 2015; Peters et al., 2010). If hybrids establish on hosts outside the narrow range of *E. crouaniorum*, opportunities for backcrossing into this species would be reduced, biasing introgression toward the more generalist *E. siliculosus* (Couceiro et al., 2015). Such ecologically mediated postzygotic isolation, where host differences constrain backcrossing, has been documented in other systems (Egan & Funk, 2009; Nosil et al., 2009), although this remains to be tested directly in *Ectocarpus*.

Secondly, differences in reproductive mode may further reinforce this asymmetry. Previous study showed that although both *Ectocarpus* species are capable of clonal reproduction in the wild, *E. crouaniorum* populations showed significantly higher levels of selfing (Couceiro et al., 2015; Montecinos, Guillemin, et al., 2017) while clonal reproduction could be dominant in some *E. siliculosus* populations (Couceiro et al., 2015). In hybridizing species pairs, a higher propensity for asexual reproduction reduces the pool of gametes available for interspecific fusion. Consequently, the more asexual species can act as a stronger barrier to incoming gene flow, while still contributing alleles to the more sexually reproducing species, whose gametes are more available for heterospecific encounters and subsequent backcrossing. This mechanism parallels the well-established effect of selfing rate on introgression directionality in flowering plants. Across 133 species pairs, (Pickup et al., 2019) showed that gene flow is typically biased from the more selfing into the more outcrossing species, a pattern also demonstrated experimentally in *Clarkia xantiana* (Sianta et al., 2024) and brown algae *Fucus* (Almeida et al., 2022; Engel et al., 2005). More generally, any process that reduces the availability of gametes for interspecific mating, whether selfing, parthenogenesis, or asexual reproduction, can generate asymmetric introgression. Whether differences in parthenogenetic propensity contribute to the observed asymmetry between *E. crouaniorum* and *E. siliculosus* remains to be explored further, and represents a promising avenue for future research.

### Role of the haploid–diploid life cycle in shaping reproductive isolation

The haploid-diploid life cycle of *Ectocarpus* may fundamentally alter the dynamics of reproductive isolation in ways that are not captured by classical diploid models of speciation. In the diploid sporophyte phase, F1 hybrids are buffered from incompatibility: heterozygosity can mask recessive Dobzhansky-Muller interactions (Turelli & Orr, 1995), and the F1 sporophyte can appear fully viable and vigorous. Selection against hybrid genotypes is instead delayed until meiosis, when recombinant haploid genotypes are exposed to selection during the gametophyte phase (Otto et al., 2015). This gametophytic phase is more efficient at purging incompatible allele combinations than zygotic selection in diploids, because every locus is hemizygous and recessive incompatibilities cannot hide (Schneider et al., 2016). Thus, efficient purging of deleterious recessive alleles in the gametophyte phase may reduce selective pressure for reinforcement compared to diploids. Moreover, in classical diploid models, the production of unfit diploid F2 and backcross hybrids in sympatry imposes a fitness cost on parents, selecting for the evolution of prezygotic barriers (Servedio & Saetre, 2003). In contrast, in haploid-diploid organisms with effective gametophytic selection, F1 sporophyte hybrids may produce few or no viable advanced-generation offspring, not only because F2 zygotes are inviable, but also because the incompatible recombinant gametophytes fail to produce functional gametes. Consequently, hybrids may impose little demographic or fitness cost at the population level: sporophyte F1s form readily but fail to generate viable diploid F2 or backcross offspring, limiting introgression without requiring strong prezygotic isolation, unless they generate direct fitness costs that impose selection for reinforcement. In this context, the distinction between pre- and postzygotic barriers becomes blurred. Gametophytic incompatibilities arise after zygote formation but effectively prevent the production of the next sporophytic generation, functioning analogously to a prezygotic barrier at the population scale. This “gametophytic postzygotic” barrier may explain the coexistence of frequent sporophyte F1 hybrids with strong overall reproductive isolation in our system.

## Supporting information

Sumplemental Tables

## Acknowledgements

We would like to thank the Roscoff Bioinformatics platform ABiMS (http://abims.sb-roscoff.fr), part of the Institut Français de Bioinformatique (ANR-11-INBS-0013) and BioGenouest network, for providing computational resources. This work was also supported by the de.NBI Cloud within the German Network for Bioinformatics Infrastructure (de.NBI) and ELIXIR-DE (Forschungszentrum Jülich and W-de.NBI-001, W-de.NBI-004, W-de.NBI-008, W-de.NBI-010, W-de.NBI-013, W-de.NBI-014, W-de.NBI-016, W-de.NBI-022). We are grateful to the Roscoff Culture Collection (RCC) for culture-room facilities. This project was supported by the ERC (grant number 864038 to S.M.C.); core funding provided by the Max-Planck-Gesellschaft, and the CNRS and Sorbonne University. L.R. was supported by a postdoctoral fellowship grant "ISORESPECTO" from the Conseil Départemental du Finistère (2021). ML was supported by Agencia Nacional de Investigación y Desarrollo, FONDECYT 1221477 and Iniciativa Cientifica Milenio grant no. NCN2024_03.

## Data Accessibility and Benefit-Sharing

Sequencing data generated in this project have been deposited in the SRA database under BioProject PRJNA1451301.The accession numbers for the raw sequence data are provided in Table S1. The VCF supporting the findings of this study is available at 10.5281/zenodo.19565816.

## Author Contributions

Conceptualization: A.P.L., M.V., C.M., L.R.; methodology: C.M., L.R., Y.F.C., M.K., F.B.H.; investigation: C.M., L.R., A.P.L.; visualization: C.M, L.R, A.P.L.; resources: S.M.C., L.R., Y.F.C, M.V.; biological material collection and culture: A.F.P., C.D., M.L.G., J.C., R.L., L.R.; data curation: C.M., L.R.; writing – original draft: A.P.L., C.M., L.R.; writing – review and editing: C.M., L.R., Y.F.C, M.K., S.M.C., A.P., M.L.G., M.V., C.D. and A.P.L.;

**Figure S1.** Choice of parameters for genotype imputation using STITCH. A) The parameter K, representing the number of ancestral haplotypes, was optimized by testing eight individuals with higher sequencing depth (5.8-8.3x) representing both species and potential hybrids. These individuals were down-sampled with Picard to reach an average depth of 0.3X, 1X, 3X, and 5X to mimic the lowest coverages in the dataset. Each dot corresponds to the average of all individuals and is colored according to the depth. Genotype imputation was then performed with STITCH on a subset of chromosomes (1–4), testing different values of *K* (5, 10, 20, 30, and 40). Performance was assessed by comparing genotype concordance (imputed genotype being equal to the hidden genotype) and calling rate (percentage of retrieved genotypes). These metrics were calculated using SNPs filtered with IS > 0.4, AD > 5, and quality ≥ 40. B) Distribution of the imputation info score (IS) used to define a filtering threshold. Shown here is an example for *K* = 30 and *ngen* = 200 across the four subset of chromosomes used for parameter optimization.

**Figure S2.** Neighbor-Joining tree of mitochondrial genomes. The tree depicts the relationships among some representative individuals from this study (marked in colored circles), with 12 recognized species of *Ectocarpus* (Denoeud et al., 2024) including *Ectocarpus sp7* (the reference genome species) and *Pilayella littoralis* as outgroups. Circles are coloured according to genomic clusters: *E. siliculosus* (blue), *E. crouaniorum* (shades of red), putative undescribed species (magenta).

**Figure S3.** Simulated individuals were derived from pure Ecro1–Esil and Ecro3–Esil lineages in Chile and Europe, respectively and 2,291 species-specific SNPs. Ten individuals were generated per hybrid class and visualized in a triangle plot. The axis of the hybrid index is relative to the *E. siliculosus* species. Only one direction of backcrossing is shown for clarity, as the reciprocal is symmetric.

**Figure S4.** A) Principal component analysis (PCA), showing cryptic lineage differentiation within the *Ectocarpus* complex in Europe (n=73). Individuals are coloured according to the NGSadmix and global PCA results; *E. siliculosus* in blue, *E. crouaniorum* in red, putative hybrids in green and the unknown *Ectocarpus* species in magenta. B) Genetic structure based on NGSadmix analysis (K=3, n=73) in Europe showing the two parental species (*E. crouaniorum* in red and *E. siliculosus* in blue) and the unknown *Ectocarpus* species in magenta.

**Figure S5.** Correlation between F_ST_ and D_XY_ across DILS locus-specific models for the following comparisons: (A) Ecro1-Esil (Europe), (B) Ecro2-Esil (Chile), (C) Ecro3-Esil (Europe) and (D) Ecro1-Esil (Chile). Loci are classified as migration-associated and isolation-associated using a posterior probability threshold of 0.7.

**Figure S6.** Circular representation of the location of barriers and the polarity of diagnostic markers for all samples (n=137) across chromosomes. The inner ring shows the location of each chromosome in alternating grey and white. Moving outwards, the next ring indicates Esil parental lineage (in blue), then the admixed Esil (light blue), the F1-type hybrids (dark green), the F1-like (light green), and Ecro parental lineage (in red). Each ring shows the genotypes of each sample at each diagnostic marker (p>0.6). Red bars are diagnostic of Ecro, blue bars are diagnostic of Esil, and green bars are heterozygous for markers diagnostic of each species. The location of each 50 kb of sequence for each chromosome is indicated on the outside of the circle.

**Table S1.** Samples used in the study.

**Table S2.** Sex markers used in the study.

**Table S3.** Posterior probabilities from NewHybrids supporting the assignment of each individual to parental species or hybrid classes.

**Table S4.** Phasing results obtained with HapCut2.

**Table S5.** Prior values for DILS analysis.

**Table S6.** Best demographic model assessed with DILS with posterior probabilities.

**Table S7.** Classification of loci identified by DILS into isolation and migration categories.

