## Supplementary figures and images for "Haplotype tagging sheds light on speciation between two divergent cryptic species of the brown algae *Ectocarpus*"

### FigS1

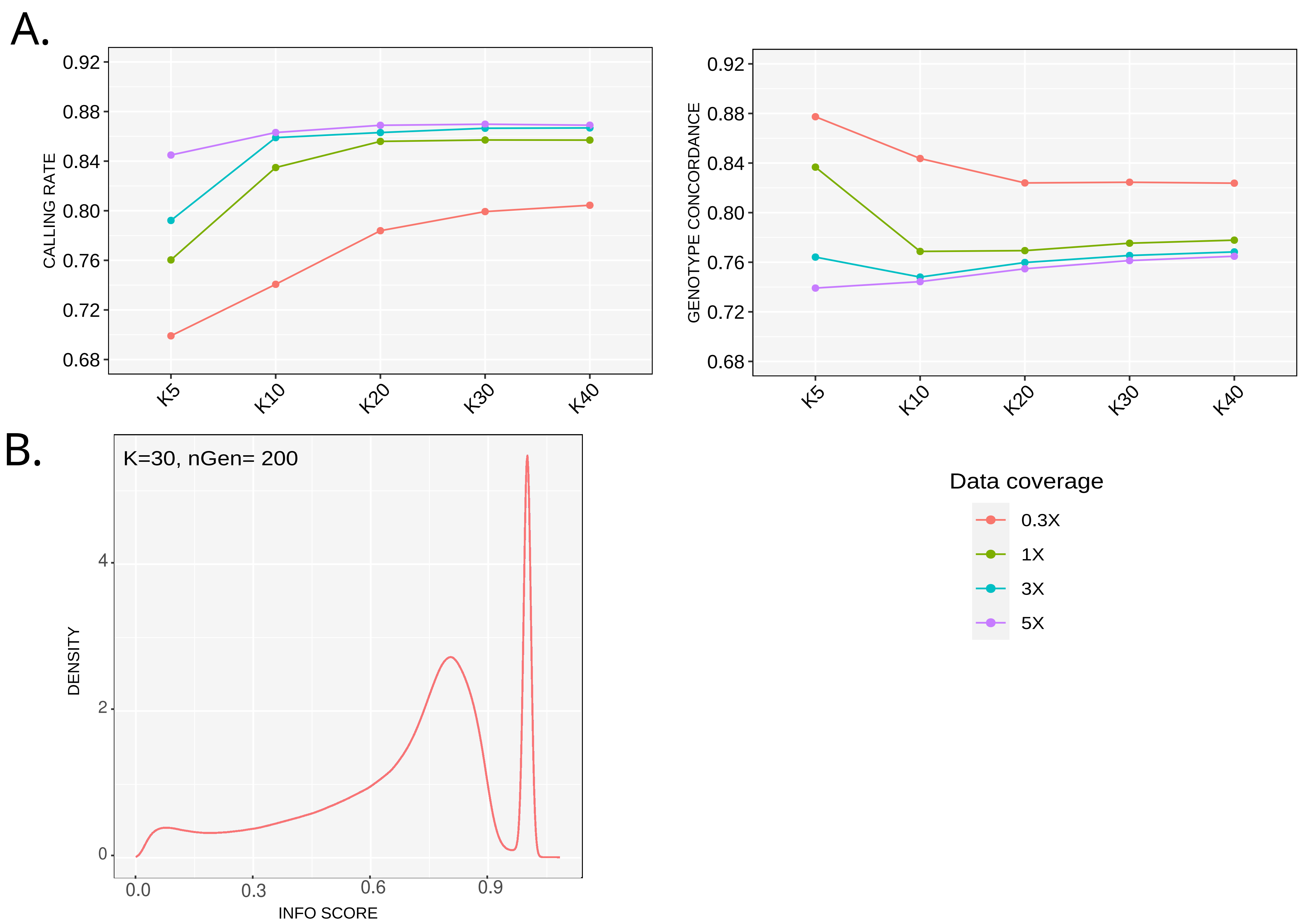

### FigS2

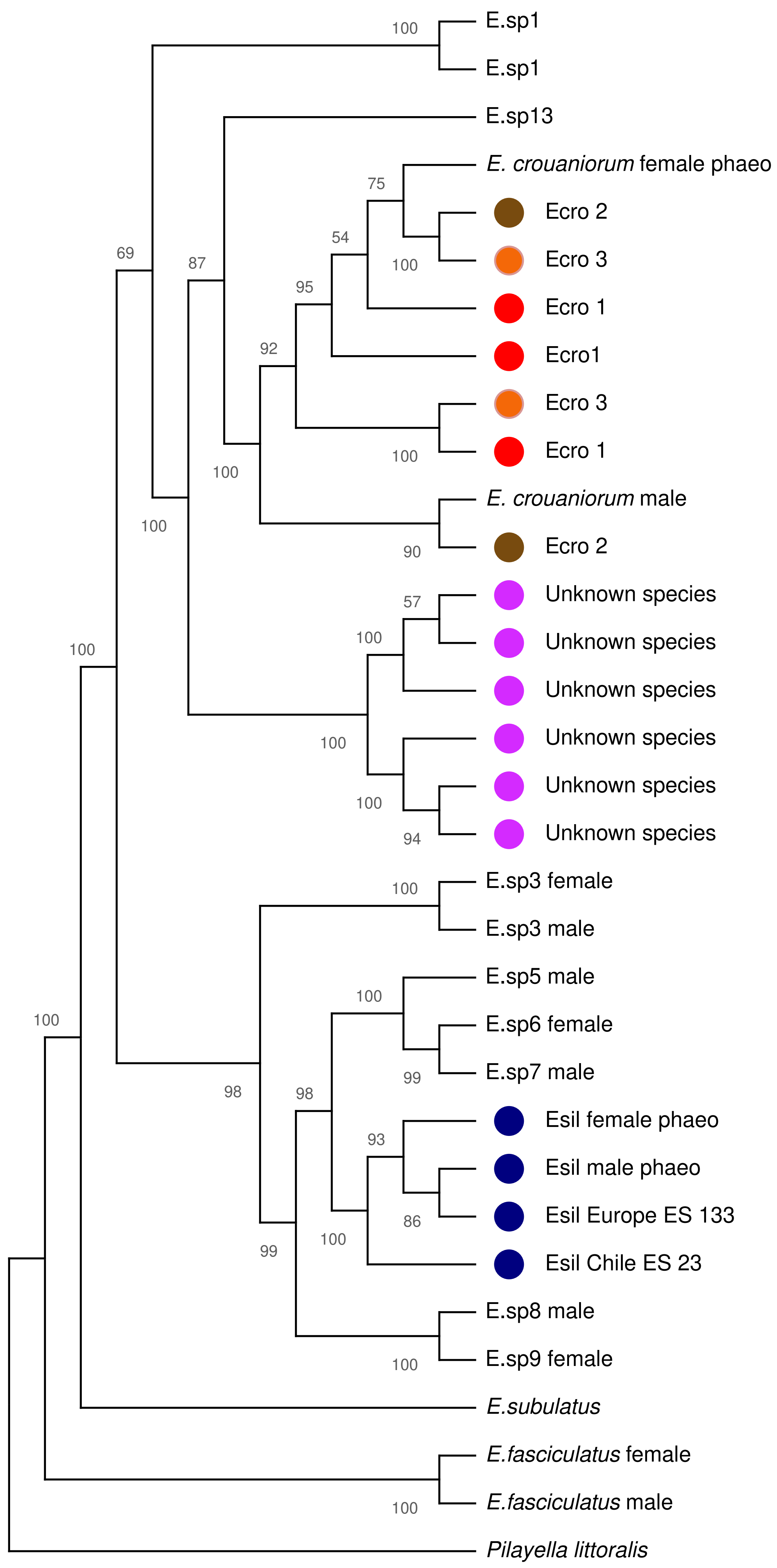

### FigS3

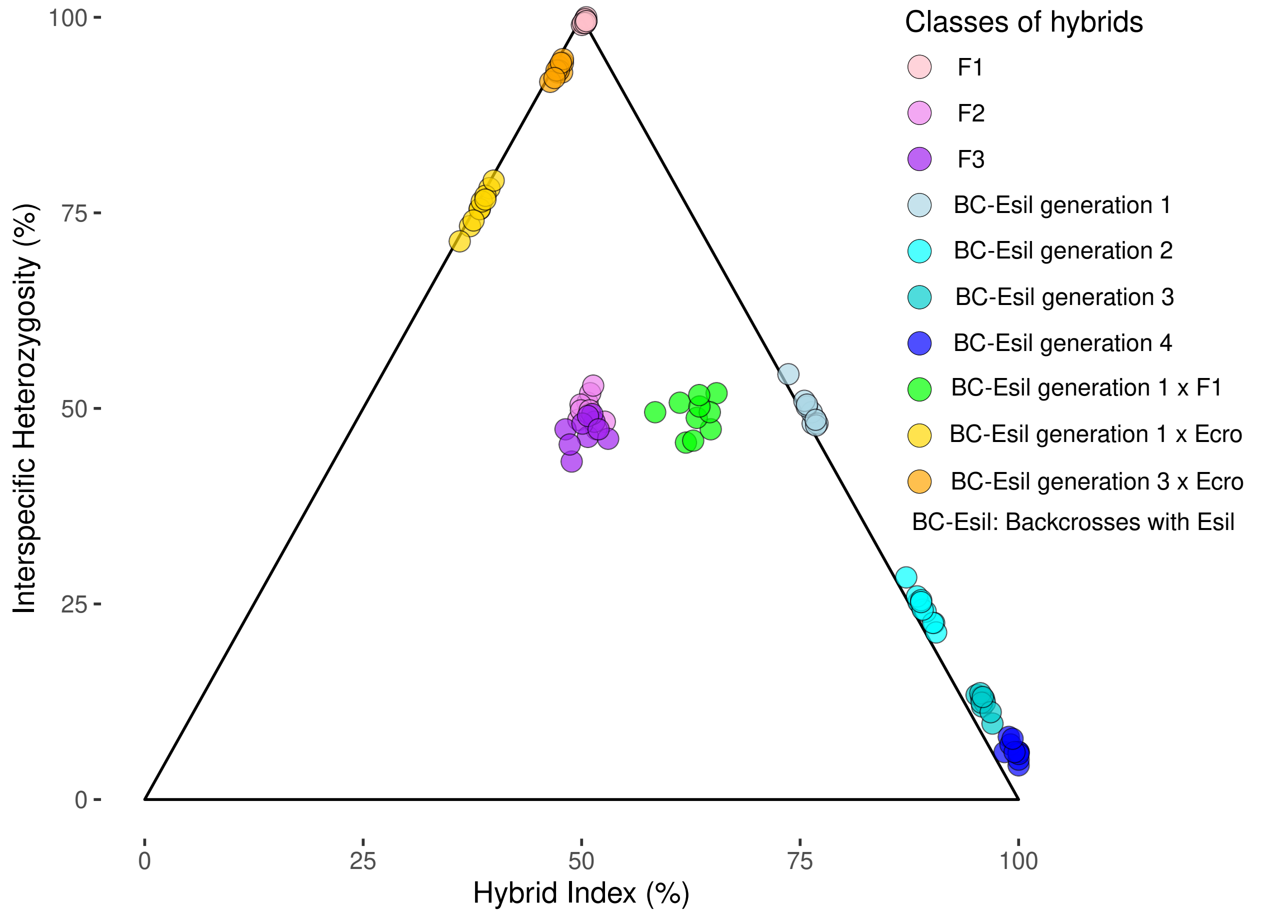

### FigS4

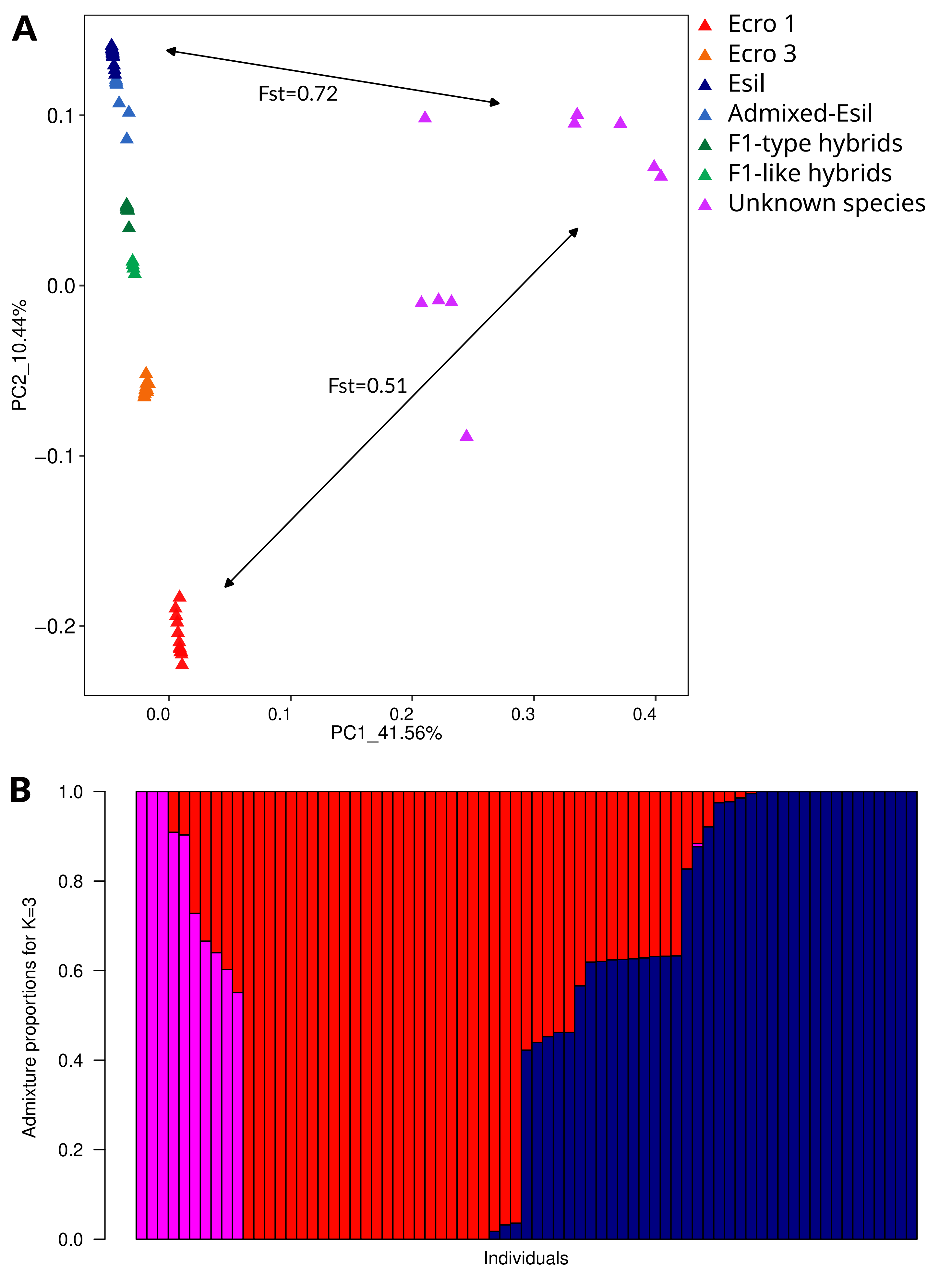

### FigS5

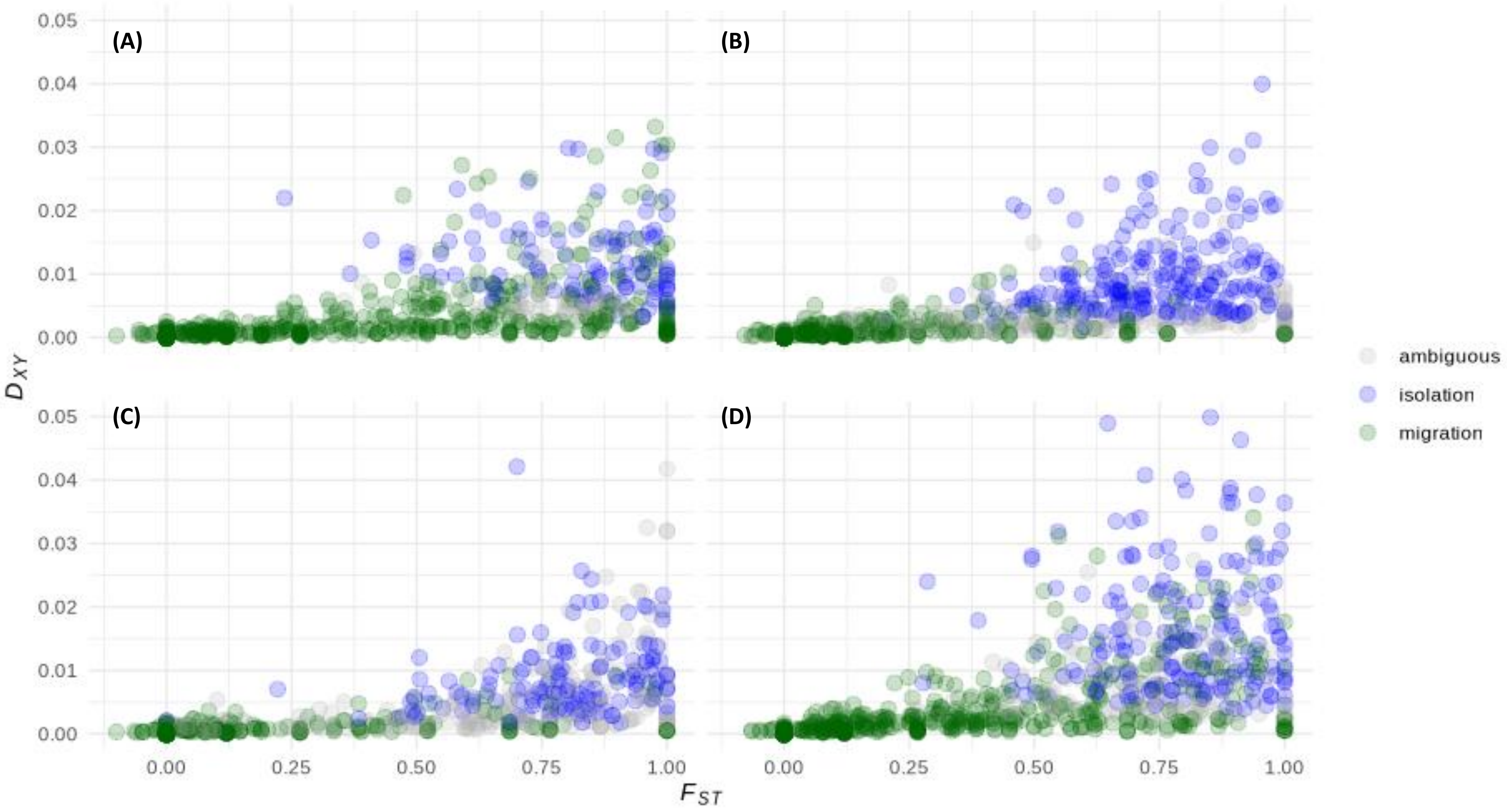

### FigS6

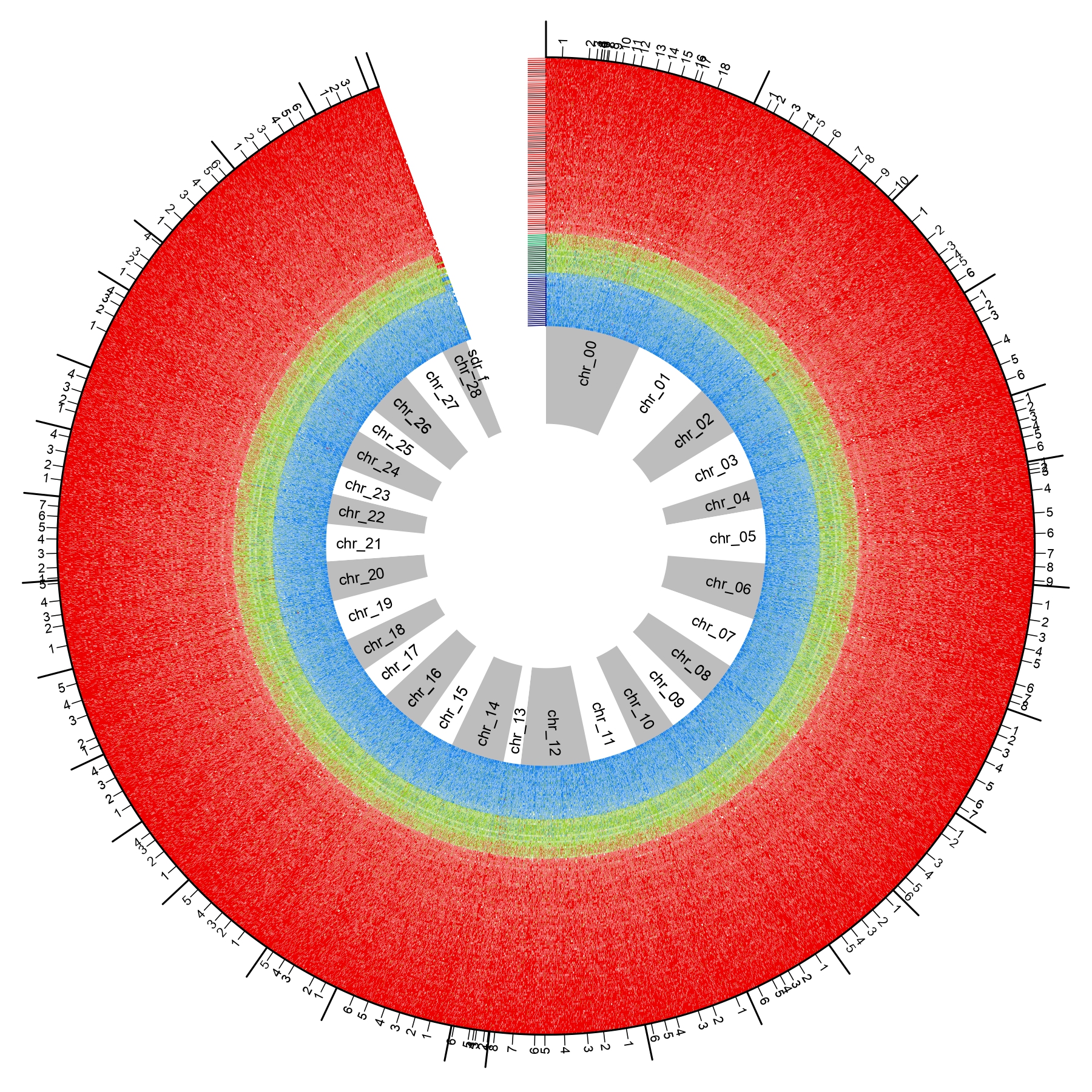

### FigS7

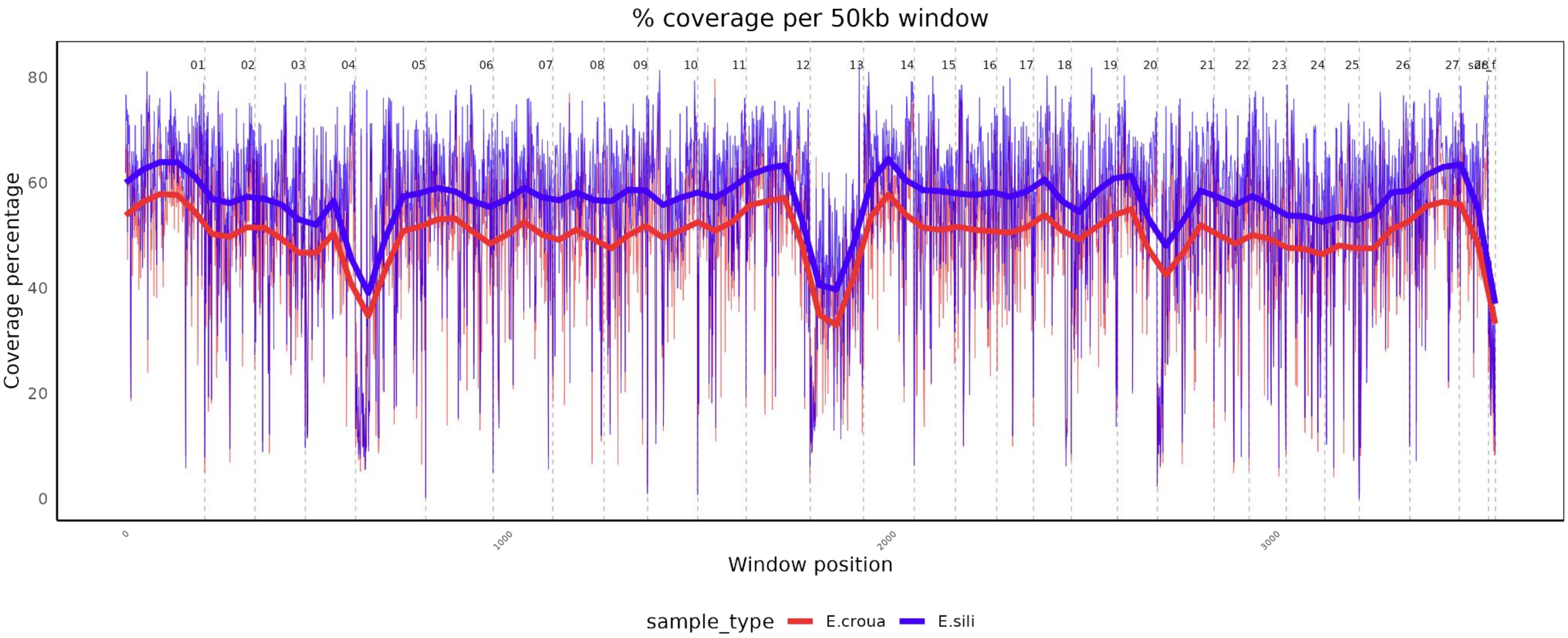
